# Mapping the Metalloproteome of *Deinococcus indicus* DR1 through Integrative Structure and Function Annotation

**DOI:** 10.1101/2025.07.11.664487

**Authors:** Sweety Deena Ramesh, Giri Vasan, Shricharan Senthilkumar, Menaka Thambiraja, Deepa Sethi, Richa Priyadarshini, Ragothaman M. Yennamalli

## Abstract

*Deinococcus indicus* DR1 is a rod-shaped bacterium isolated from the Dadri wetlands (Uttar Pradesh, India) that tolerates ionizing radiation and arsenic. The molecular basis of its wider heavy-metal resilience, particularly among the 1017 out of 4128 proteins still annotated as hypothetical, remains unclear. We performed a proteome-wide structural and functional survey to address this gap. All the 4128 proteins were modeled with AlphaFold2, yielding very-high-confidence structures (pLDDT ≥ 90) for 2145 sequences. CATH/InterPro analysis assigned domains to 2735 proteins. Functions were predicted by combining DeepFRI (graph neural-network), MorphologFinder (Foldseek plus EggNOG-Mapper), and existing GenBank annotations. The integrated workflow suggests that more than 100 previously uncharacterized proteins may bind or transport arsenic, chromium, cobalt, copper, iron, manganese, molybdenum, nickel, or zinc, indicating a metal-handling capacity that extends beyond the known ars operon. Recurrent domain architectures that include P-loop NTPases, Rossmann folds, GNAT acetyl-transferases, and sensor modules (CHASE, PAS, GAF) point to coordinated redox regulation and efflux pathways. Twenty high-confidence metal-binding candidates have been prioritized for experimental validation through expression, mutagenesis, and knockout studies. All structural models, domain assignments, and query tools are available at https://deinococcus.in, providing a resource for future investigations of heavy-metal tolerance in this organism.

## 1. Introduction

Heavy metals are naturally occurring elements, but their uncontrolled release from volcanic eruptions, mining, smelting, coal combustion, and industrial wastewater has pushed their concentrations far beyond background levels in many ecosystems. Chronic exposure to metals such as arsenic, cadmium, and chromium is associated with cancers, immune dysfunction, and disruption of the gut microbiota (Eggers et al., 2018). Microorganisms in contaminated niches face intense selective pressure and have consequently evolved an impressive repertoire of metal-detoxification strategies (Igiri et al., 2018). These include extracellular sequestration, intracellular chelation, enzymatic redox cycling that converts a toxic species to a less harmful form, and active export through ATP-driven or proton-motive pumps. Genomic surveys illustrate the breadth of this adaptation. *Escherichia coli* E308, for example, carries entire cassettes of resistance genes: arsB, arsC, and arsR for arsenic; copC, copD, copG, cusA, cusB, cusC, cusF and cusR gene cluster for copper; rcnA, rcnB, rcnC, nikA, nikB, nikC, nikD, nikE and nikR gene cluster for nickel; rcnA, rcnB, rcnC and corC for cobalt; mgtC for magnesium; mntH for manganese; silE for silver; tehA and tehB for tellurium; and zntA and zntB for Zinc (Yang et al., 2020; Sharma et al., 2000). Likewise, *Staphylococcus aureu*s deploys CopB and Mco for copper, CadA/CadB for cadmium, and czrC for dual zinc– cadmium defense (Tsai et al., 1992; Cavaco et al., 2010). These examples show that metal stress shapes microbial genomes.

Among extremophiles, the family Deinococcaceae stands out for its unparalleled resistance to ionizing radiation, desiccation, UV, and oxidative bursts. Such cross-protection is rooted in superb DNA-repair networks, manganese-based antioxidant systems, and carotenoid shielding pigments. Many Deinococcus species also tolerate—or even immobilize—heavy metals through biosorption and biomineralization (Jaafar et al., 2016). Over sixty members have been recovered from environments as varied as desert sand, Antarctic ice, hot springs, radioactive waste, spacecraft surfaces, and arsenic-laden aquifers (Gerber et al., 2015; Weon et al., 2007; Ghosh et al., 2010; Chauhan et al., 2019). Their cosmopolitan distribution implies robust, transferable solutions to multiple stressors.

*Deinococcus indicus* was first isolated from arsenic-contaminated pond water in West Bengal, India. The type strain is a non-motile, red-pigmented rod-shaped that withstands intense UV and tolerates arsenic up to millimolar concentrations (Suresh et al., 2004; Chauhan et al., 2017; Dhanapal et al., 2021). A second isolate, *D. indicus* DR1, was recovered from the Dadri wetland in Uttar Pradesh and characterized as an aerobic, spore-forming chemo-organotroph that produces the carotenoid deinoxanthin, accounting for its pink hue (Ranganathan et al., 2023). *D. indicus* DR1 grows from 4 °C to 55 °C, a temperature span rare even within extremophiles, and reduces arsenate [As (V)] to the more mobile arsenite [As (III)], which it then expels via an ArsB efflux pump (Chauhan et al., 2019).

Arsenic resistance in *D. indicus* DR1 is encoded by a six-gene ars cluster comprising two reductases (ArsC2, ArsC3), two transcriptional repressors (ArsR1, ArsR2), a metallophosphatase, and the membrane transporter ArsB. The reductases convert arsenate to arsenite, the transcription factors modulate operon expression in response to metal load, and ArsB pumps arsenite out of the cytosol. Comparative genomics shows remarkable diversity in ars architecture: the R773 plasmid of *E. coli* carries a five-gene cassette (arsR, D, A, B, C), whereas *Staphylococcus aureus* limits itself to arsR, B, and C, and Herminiimonas arsenicoxydans harbors four distinct ars operons, reflecting its ancient adaptation to arsenic-rich geology (Ranganathan et al., 2023). *D. indicus* DR1 thus provides an ideal model to explore how a radiation-hardy background couples with metal detoxification.

Yet the *D. indicus* DR1 genome, like most bacterial genomes, contains a large fraction of hypothetical proteins (HPs)—open reading frames with no experimentally determined function. Such sequences may encode novel enzymes, regulators, or transporters that have escaped annotation because they lack homology to characterized proteins or contain domains of unknown function (DUFs) (Ijaq et al., 2019). DUFs alone account for thousands of structures in the Protein Data Bank, underscoring how often structural genomics outpaces functional assignment (Bharat Siva Varma et al., 2015). Because heavy-metal tolerance is a multigenic trait, uncharacterized proteins could hold the key to unexplained phenotypes.

Domains are the evolutionary building blocks of proteins; they combine in modular fashion to create complex activities (Aziz and Caetano-Anollés, 2021). Domain-centric analysis has clarified multidrug resistance in fungi, magnetosome formation in bacteria, and host adaptation in viruses. For instance, Barrera et al. (2014) screened 137 fungal proteomes for domains linked to lovastatin biosynthesis and siroheme pathways, pinpointing new antifungal targets. Applying a similar lens to *D. indicus* DR1 would uncover domain arrangements unique to arsenic defense—information that can guide biochemical tests or metabolic engineering.

The overarching goal of this work is to annotate DR1’s hypothetical proteins and classify the structural domains of its 4128-member proteome to reveal the molecular underpinnings of its heavy-metal resistance. Structural characterization will map arsenic-resistance domains, while functional annotation of HPs will identify novel contributors to DR1’s exceptional tolerance. By illuminating the structural basis of heavy-metal resistance in *D. indicus* DR1, our work addresses both fundamental and applied questions. On the basic side, it enriches our understanding of how extremophiles coordinate multiple stress responses—radiation, desiccation, and metal toxicity—within a single cellular framework. On the applied side, new reductases or transporters from DR1 could inform bio-remediation strategies aimed at detoxifying arsenic-contaminated water, a pressing public-health issue in South and Southeast Asia. Moreover, elucidating domain architectures expands the catalog of protein scaffolds available for synthetic biology, nanomaterial binding, and biosensor design.

## 2. Materials and Methods

### 2.1 Structure prediction

Accurate structural models are essential for probing protein function at atomic resolution. To characterize the *D. indicus* DR1 proteome, we followed the workflow outlined in Figure 1. The complete set of protein sequences (FASTA) was downloaded from NCBI (assembly accession GCA_002198095.1). The data set contains 4,128 proteins, of which 1,017 are annotated as hypothetical—i.e., no function has yet been assigned in GenBank. All sequences were modeled with AlphaFold2, whose deep-learning framework routinely yields near–experimental accuracy when only the amino-acid sequence is provided (Jumper et al., 2021; Varadi et al., 2022).

**Figure 1:**
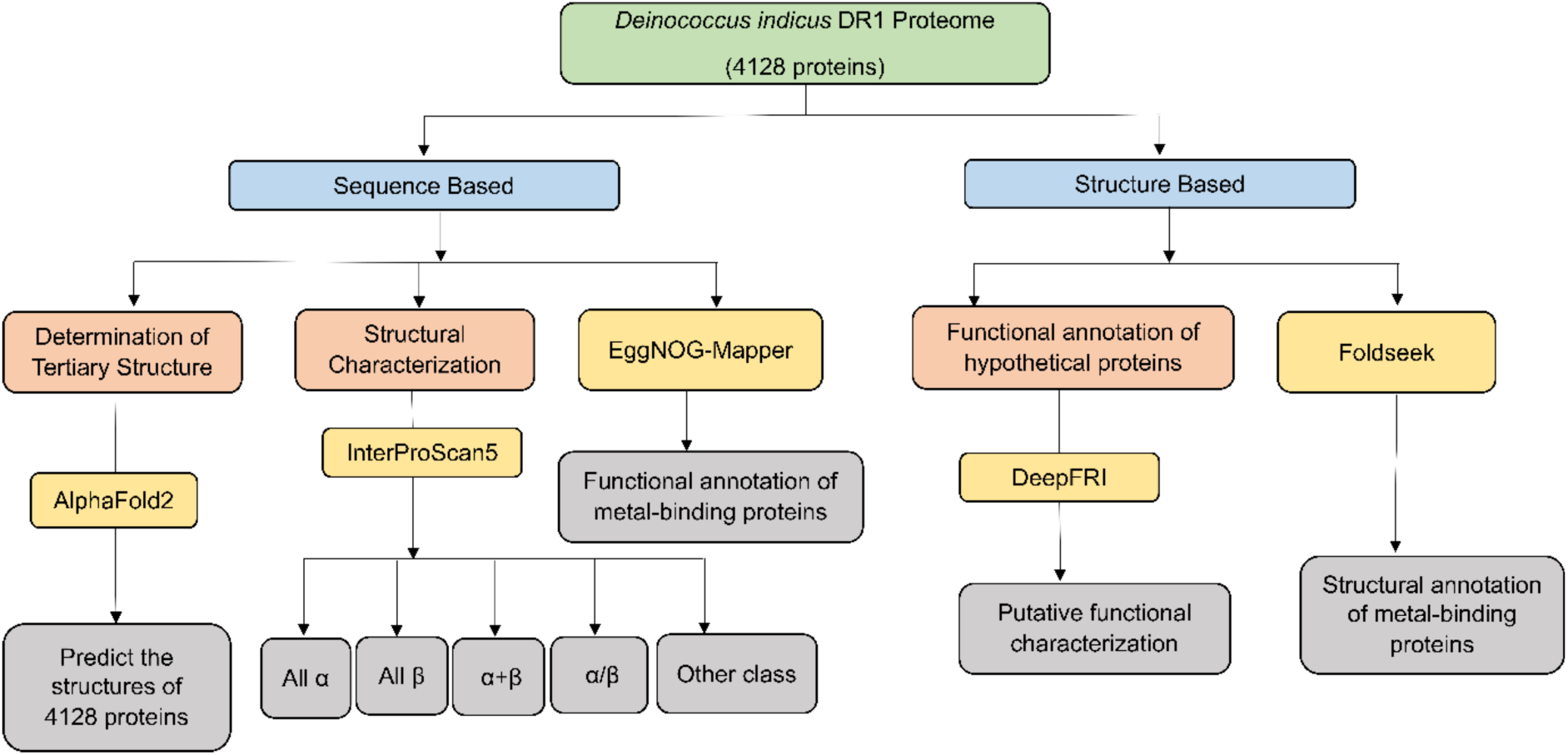
Workflow of structure prediction and functional annotation of *D. indicus* DR1 proteome. Structural characterization using the sequences of *D. indicus* DR1 gives the CATH classification of the proteins. Functional annotation and clustering were done using the structures that have been predicted using AlphaFold2.

### 2.2 Structural characterization

Protein domains are compact, evolutionarily conserved units that often dictate molecular function. Classifying these domains across the *D. indicus* DR1 proteome therefore provides a direct route to discover motifs linked to arsenic resistance. We carried out domain annotation with InterProScan5, a standalone Java package widely used by UniProtKB and genome-sequencing pipelines to generate first-pass functional profiles (Jones et al., 2014).

InterProScan accepts primary sequences and searches them against multiple profile and HMM libraries—including Pfam, TIGRFAM, SMART, PIRSF, PANTHER, HAMAP, PROSITE, ProDom, PRINTS, CATH-Gene3D, and SUPERFAMILY—using algorithms such as BLAST and HMMER. It also integrates TMHMM, SignalP, and Phobius to flag transmembrane segments and signal peptides. The software post-processes raw hits to remove redundancy and outputs results in XML, JSON, and TSV formats. For the present study we retained only the CATH-Gene3D section, which assigns each sequence to a CATH structural domain and supplies the corresponding CATH identifier. This filtered output forms the basis for downstream analyses of domain distribution and potential heavy metal tolerance mechanisms.

### 2.3 Functional annotation

Hypothetical proteins in the *D. indicus* DR1 proteome were functionally annotated with DeepFRI, a geometric deep-learning framework that combines protein language models with graph convolutional networks (GCNs) to predict Gene Ontology (GO) terms (Gligorijević et al., 2021). The GO consortium organizes protein function hierarchically into Enzyme Commission (EC) numbers and three major ontologies: Molecular Function (MF), Biological Process (BP), and Cellular Component (CC). Traditional sequence-based methods often struggle with remote homology; GCNs mitigate this limitation by encoding each protein as a graph in which residues are nodes and spatial contacts are edges, allowing the network to learn structure-aware features.

DeepFRI operates in two stages. First, a pre-trained, task-agnostic transformer generates a rich sequence embedding, while the atomic coordinates are converted into a residue-interaction graph. Second, the combined representation passes through a GCN that outputs probabilities for thousands of GO terms. A gradient-weighted class activation map (Grad-CAM) highlights structural regions most responsible for each prediction, providing residue-level interpretability. In this study, we supplied AlphaFold2-derived coordinates for all 1,017 hypothetical proteins; the resulting GO annotations and Grad-CAM maps guided the identification of previously uncharacterized enzymes and metal-binding factors potentially involved in arsenic tolerance.

### 2.4 Implementation of MorphologFinder workflow for metal binding and uncharacterized proteins

To obtain complementary functional clues, we applied the MorphologFinder single-protein workflow, which integrates structure- and sequence-based annotation. For each hypothetical protein we supplied (i) the AlphaFold2 model and (ii) its amino-acid sequence.

The 3-D model was submitted to the Foldseek server (https://search.foldseek.com), and structural neighbors were retrieved from the PDB, UniProt/Swiss-Prot, and AlphaFold model databases. For every database we retained the single best hit—defined as the match with the lowest E-value. Functional information associated with those top hits served as the structure-derived annotation.

The same protein sequence was analyzed with EggNOG-Mapper (http://eggnog-mapper.embl.de/), which assigns orthology-based Gene Ontology terms, KEGG pathways, and functional descriptions.

Annotations from Foldseek and EggNOG-Mapper were merged and grouped by metal keyword (arsenic, copper, zinc, etc.). We then cross-checked these assignments against the metal-binding predictions generated by DeepFRI. If MorphologFinder produced no annotation for a given protein, we substituted the corresponding GenBank description. This combined dataset enabled a focused list of candidate proteins potentially involved in heavy-metal binding and detoxification.

## 3. Results

### 3.1 Structure prediction

All 4,128 proteins from the *D. indicus* DR1 genome were modeled with AlphaFold2. For each sequence the program produces five candidate structures and assigns two built-in confidence metrics: the per-residue Predicted Local Distance Difference Test (pLDDT) and the global Predicted Template Modeling score (pTM). We retained the single model with the highest mean pLDDT for every protein.

The pLDDT ranges from 0 to 100 and is interpreted as follows: scores ≥ 90 indicate a high-confidence, near–experimental model; 70–89 denote medium confidence; 50–69 mark low confidence; and values < 50 flag very low confidence. The distribution of pLDDT values across the DR1 proteome is listed in Supplementary Table 1 and graphed in Figure 2.

**Figure 2:**
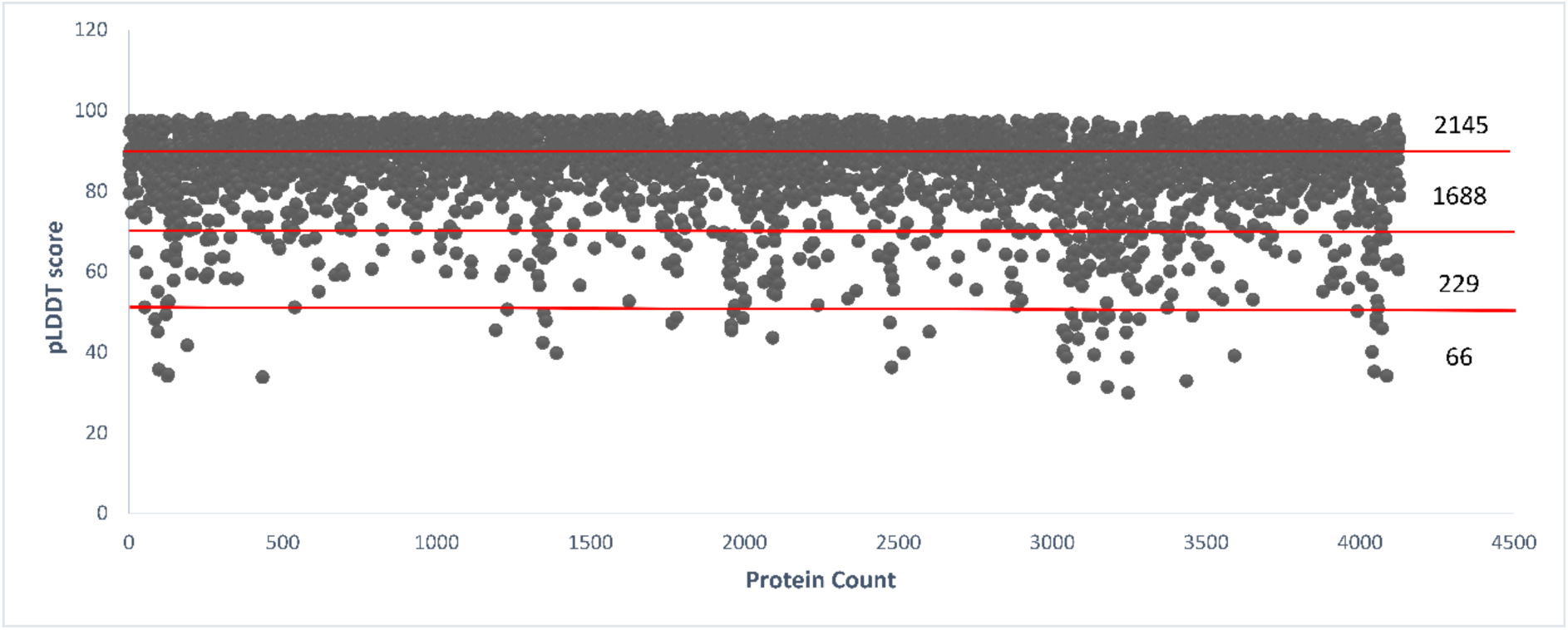
The distribution of pLDDT score among the AlphaFold2 predicted structures. The pLDDT scores of the protein structures of the D. indicus DR1 strain that was predicted using AlphaFold2. The numbers on the right indicate the number of proteins that have pLDDT scores at various cut-offs.

AlphaFold2 delivered high-confidence models (pLDDT ≥ 90) for 2145 proteins—just over half of the proteome. A further 1688 sequences fall in the medium-confidence bracket. Only 229 proteins yielded low-confidence structures (pLDDT 50–69), and 66 scored below 50. Thus, reliable atomic models are available for the vast majority of D. indicus DR1 proteins, with relatively few cases requiring additional validation.

### 3.2 Structural characterization

Protein domains are widely regarded as the basic structural, functional, and evolutionary units of proteins (Basu et al., 2009). A given polypeptide may fold into a single domain or into several domains linked into a multidomain architecture. In structural terms a domain is an independently folded unit, whereas in sequence analysis a domain is detected as a highly conserved region (Wang et al., 2021). Because domain makeup often governs molecular activity, mapping every domain in the *D. indicus* DR1 proteome provides a direct window onto the motifs that could underlie arsenic tolerance.

Domain annotation was carried out with InterProScan 5, and only the CATH-Gene3D section of the CSV output—which supplies both CATH classifications and residue ranges—was retained for analysis (Jones et al., 2014). Of the 4128 proteins, 2735 matched at least one CATH entry, whereas 1393 lacked any hit, suggesting the presence of domains not yet represented in current databases. Within the annotated set, one domain (CATH ID 3.40.50.300) occurs 272 times, several others appear more than one hundred times, and many recur more than fifty times (Figure 3; Table 1).

**Figure 3:**
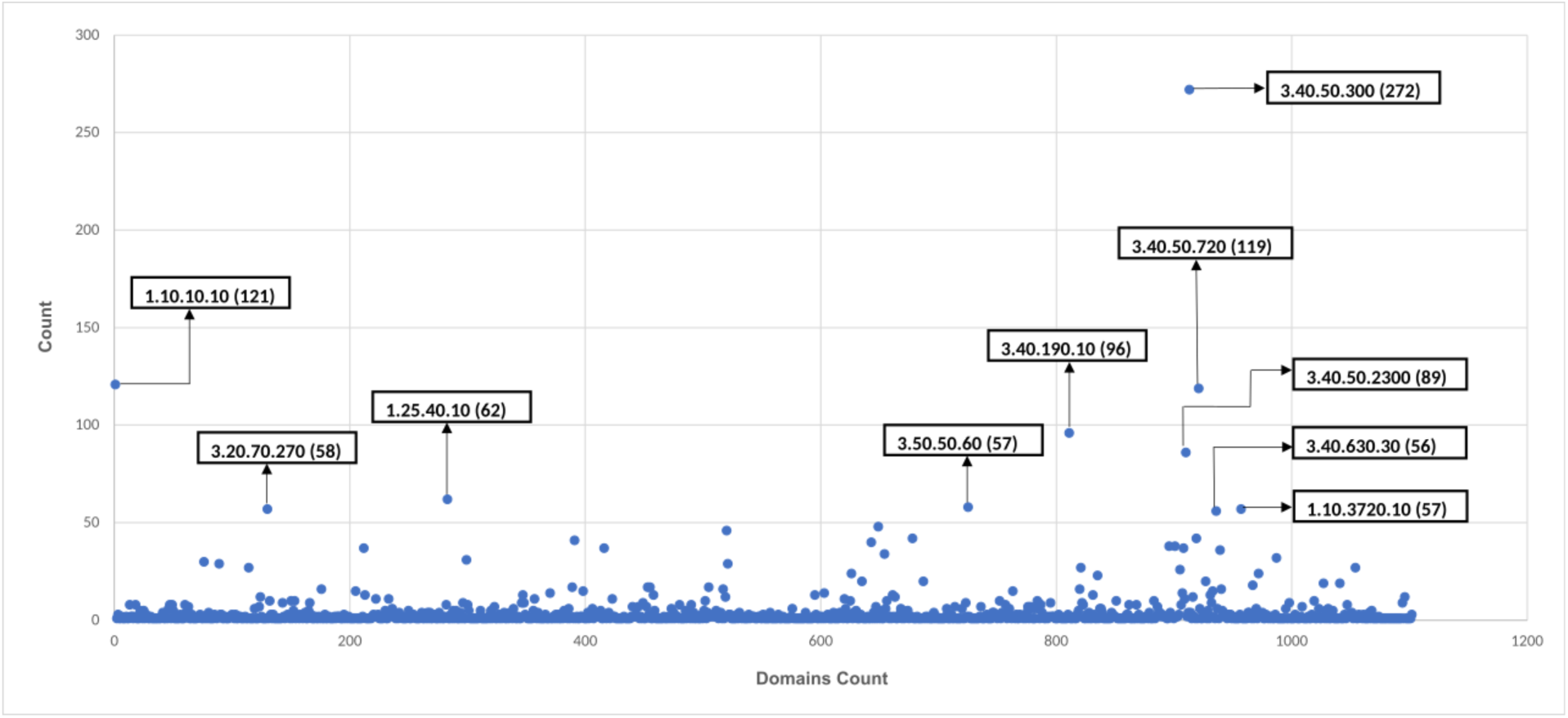
CATH classification domains of *D. indicus* DR1 proteome, showing the frequency of repeated domains. Here the domains that get repeated more than 50 times are labelled.

**Table 1:** List of highly occurring CATH classified domains. The functions of most occurred protein domains in *D. indicus* DR1 and its name.

| S. no. | CATH ID | Number of occurrences | Domain name |
| --- | --- | --- | --- |
| 1. | 3.40.50.300 | 272 | P-loop containing nucleotide triphosphate hydrolase |
| 2. | 1.10.10.10 | 121 | Winged double helix-like DNA-binding domain |
| 3. | 3.40.50.720 | 119 | NAD(P)-binding Rossmann-like domain |
| 4. | 3.40.190.10 | 96 | Periplasmic binding protein-like II |
| 5. | 3.40.50.2300 | 86 | Response regulator |
| 6. | 1.25.40.10 | 62 | Tetratricopeptide repeat domain |
| 7. | 3.30.70.270 | 58 | Reverse transcriptase |
| 8. | 3.50.50.60 | 57 | FAD/NAD(P)-binding domain |
| 9. | 1.10.3720.10 | 57 | MetI-like domain |
| 10. | 3.40.630.30 | 56 | Gcn5-related N-acetyltransferase (GNAT) |

**Figure 4:**
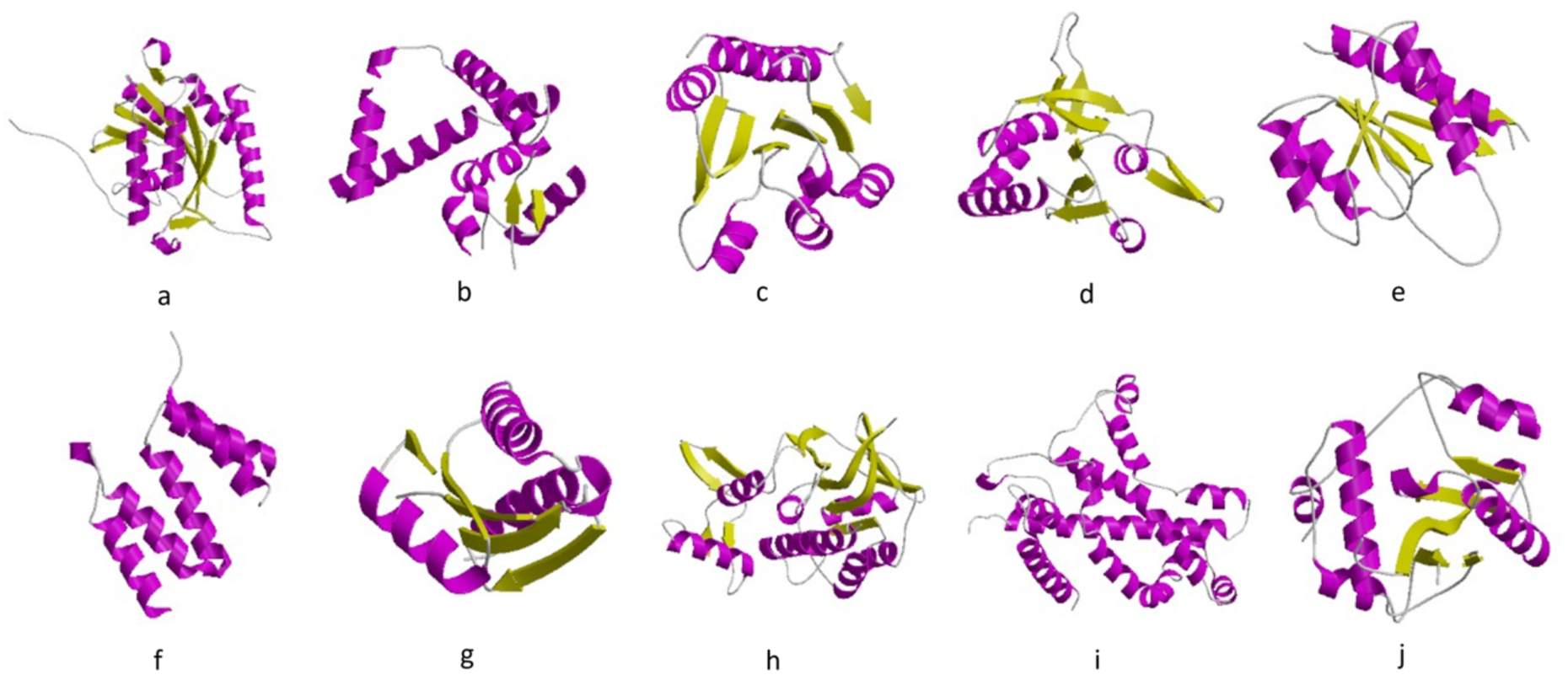
Highly repeated protein domains in the proteome of D. indicus DR1 predicted by InterProScan5. It shows the ten domains that repeated for multiple times in the proteome. (a) P-loop containing nucleotide triphosphate hydrolase. (b) Winged double helix-like DNA-binding domain. (c) NAD(P)-binding Rossmann-like domain. (d) Periplasmic binding protein-like II. (e) Response regulator (f) Tetratricopeptide repeat domain. (g) Reverse transcriptase. (h) FAD/NAD(P)-binding domain. (i) MetI-like domain. (j) Gcn5-related N-acetyltransferase (GNAT).

The proteome displays a striking range of architectural complexity. InterProScan identifies 1 553 single-domain proteins, 775 two-domain proteins, 258 three-domain proteins, 81 four-domain proteins, 44 five-domain proteins, 17 six-domain proteins, six seven-domain proteins, and a single twelve-domain protein (Figure 5; Table 2; Supplementary Table 2).

**Figure 5:**
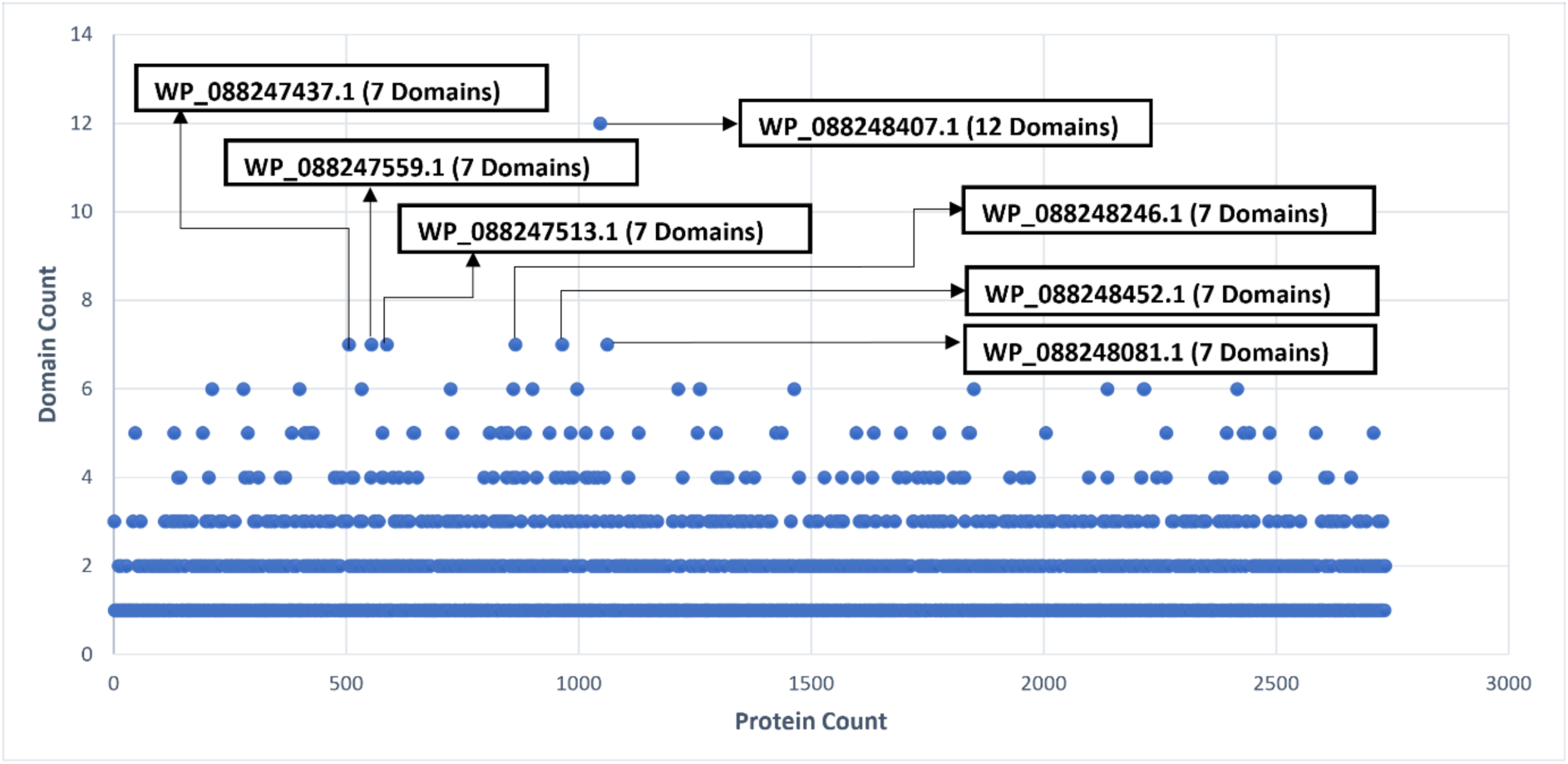
Multi-domain proteins having seven or more domains predicted by InterProScan5. It represents the number of occurrences of all the domains in the D. indicus DR1 proteome. Domains that get repeated more than seven times are labelled and the domain counts were also represented.

**Table 2:**
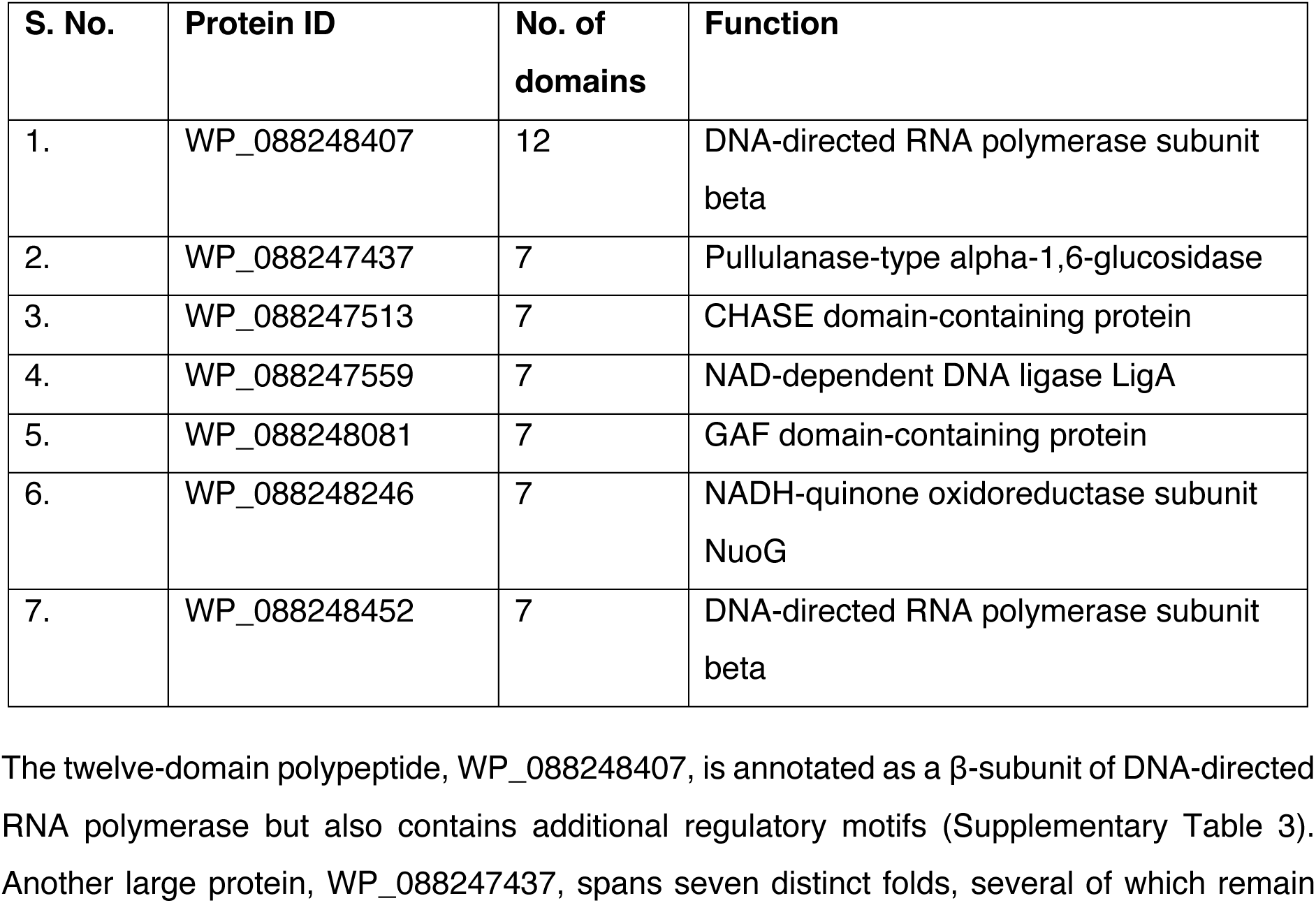
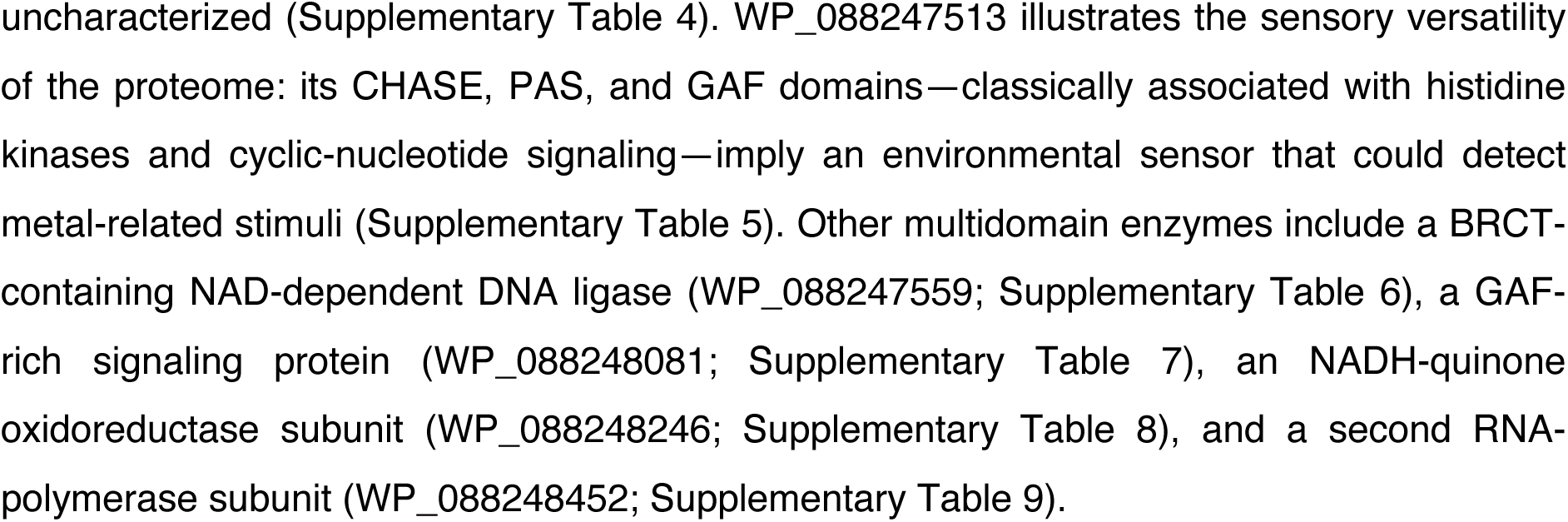
Multi-domain proteins having seven or more domains predicted by InterProScan5. Proteins of *D. indicus* DR1 with domain count of more than seven and the function of those proteins that were predicted using InterProScan5.

Taken together, the dominance of nucleic-acid-binding, redox, and sensor domains, coupled with the prevalence of elaborate multidomain arrangements, suggests that *D. indicus* DR1 deploys an extensive network of regulatory and metabolic proteins to monitor and mitigate arsenic and other heavy-metal stress.

### 3.3 Functional annotation

Deep-learning methods have transformed many areas of computational biology, and graph-based neural networks now match or exceed traditional approaches for function prediction (Bryant P, 2023) While early efforts stored protein structures as dense three-dimensional grids for 3-D convolutional neural networks—an approach that wastes memory because most voxels contain empty space—newer models treat a protein as a graph whose nodes are residues and whose edges represent spatial contacts. This representation is both compact and well suited to relational learning.

We used DeepFRI to assign functions to the 1017 hypothetical proteins in the *D. indicus* DR1 proteome. DeepFRI combines a transformer-based language model, which embeds the primary sequence, with a graph-convolutional network that encodes the residue-contact map derived from the AlphaFold2 structure (Gligorijević et al., 2021). The network outputs probabilities for Gene Ontology (GO) terms across the three major ontologies—Molecular Function (MF), Biological Process (BP), and Cellular Component (CC)—as well as Enzyme Commission (EC) numbers. A gradient-weighted class-activation map highlights the residues that drive each prediction, providing residue-level interpretability. These annotations form the basis for downstream comparisons with the CATH domain catalogue and the MorphologFinder results. Each putative function is returned with a confidence score between 0 and 1, and the highest-scoring term is taken as the primary prediction.

All 1017 hypothetical proteins from *D. indicus* DR1 were processed through DeepFRI, and the MF output was mined for GO terms containing “metal ion binding.” Table 3 lists the proteins whose metal-binding score is ≥ 0.50; five of them—WP_172417986, WP_088249605, WP_088247726, WP_172417877, and WP_172418100—show exclusively metal-related functions at high confidence.

**Table 3:**
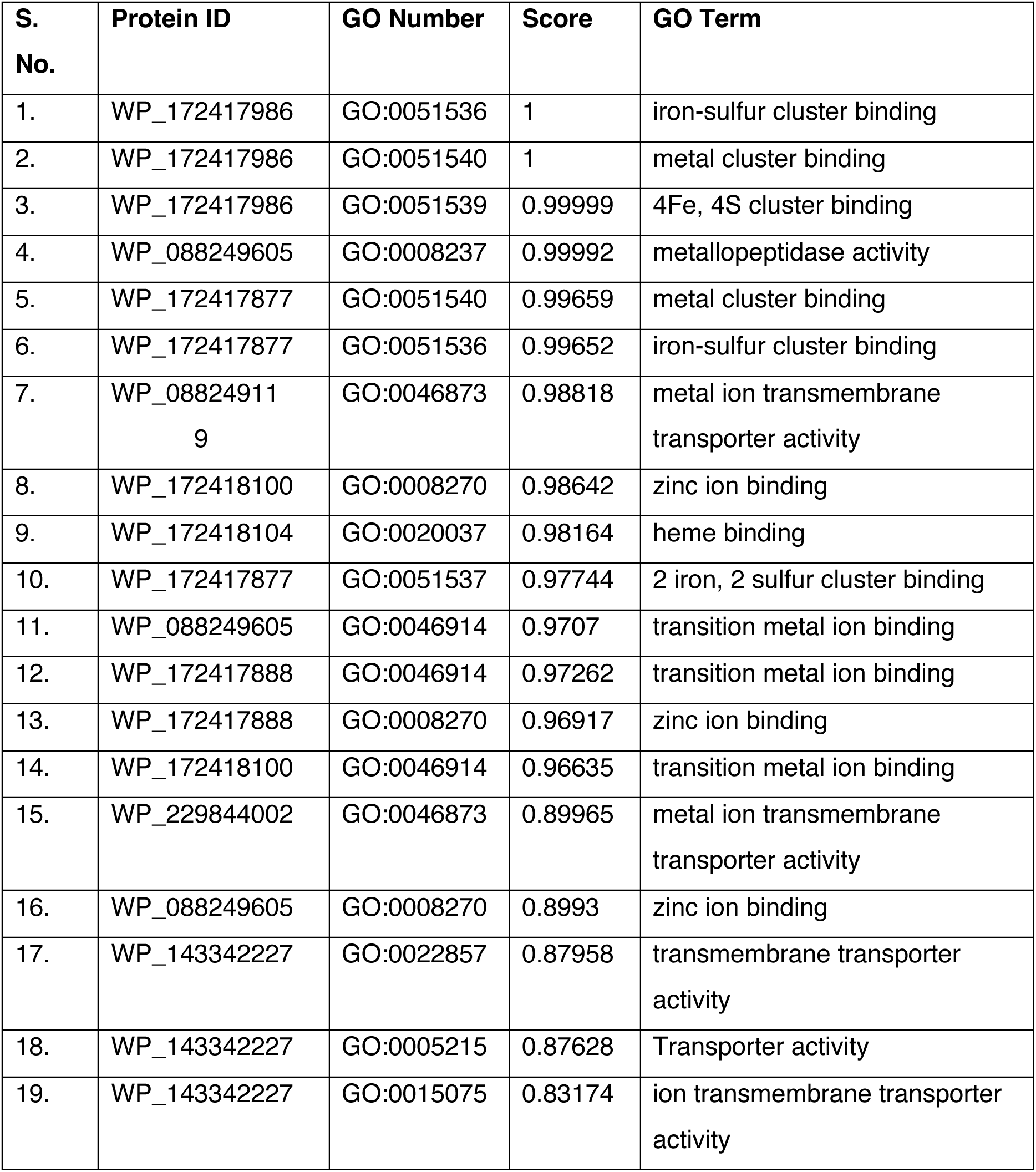
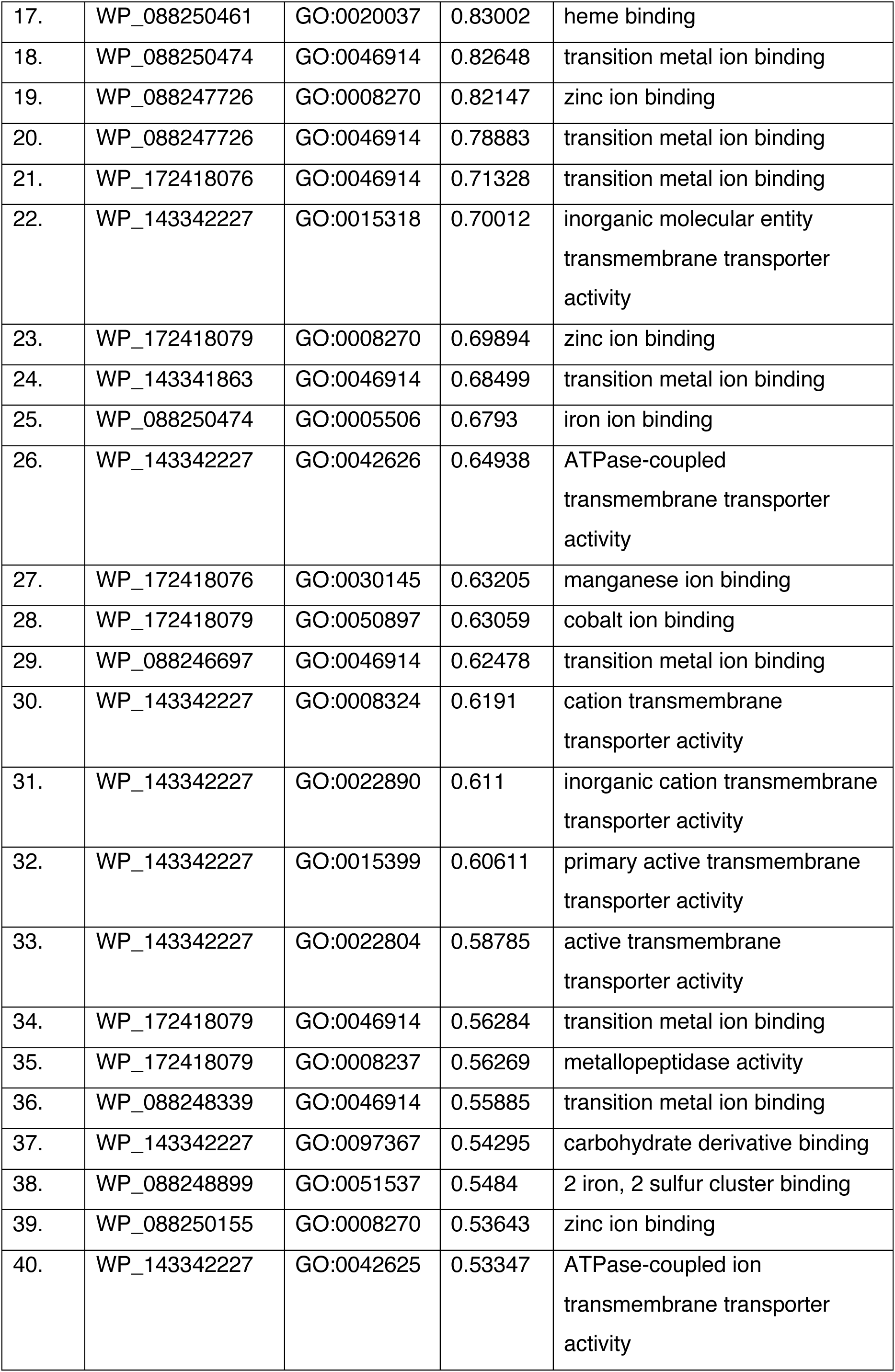
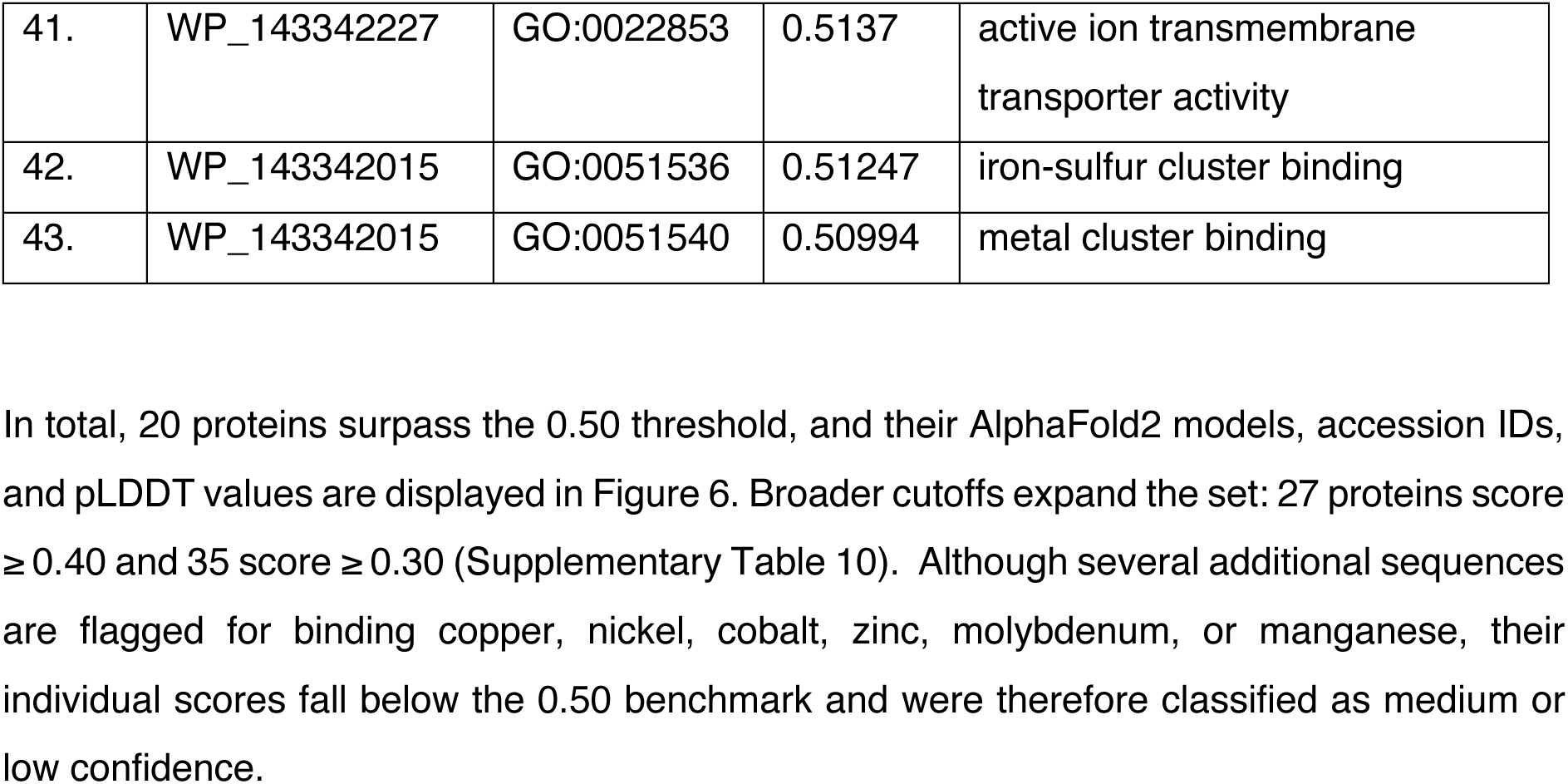
Functions of hypothetical proteins that are having a score greater than 0.5 with metal ion binding property. The annotated functions of hypothetical proteins that are having metal ion binding properties with a score of greater than 0.5.

| S. No. | Protein ID | GO Number | Score | GO Term |
| --- | --- | --- | --- | --- |
| 1. | WP_172417986 | GO:0051536 | 1 | iron-sulfur cluster binding |
| 2. | WP_172417986 | GO:0051540 | 1 | metal cluster binding |
| 3. | WP_172417986 | GO:0051539 | 0.99999 | 4Fe, 4S cluster binding |
| 4. | WP_088249605 | GO:0008237 | 0.99992 | metallopeptidase activity |
| 5. | WP_172417877 | GO:0051540 | 0.99659 | metal cluster binding |
| 6. | WP_172417877 | GO:0051536 | 0.99652 | iron-sulfur cluster binding |
| 7. | WP_08824911<br>9 | GO:0046873 | 0.98818 | metal ion transmembrane transporter activity |
| 8. | WP_172418100 | GO:0008270 | 0.98642 | zinc ion binding |
| 9. | WP_172418104 | GO:0020037 | 0.98164 | heme binding |
| 10. | WP_172417877 | GO:0051537 | 0.97744 | 2 iron, 2 sulfur cluster binding |
| 11. | WP_088249605 | GO:0046914 | 0.9707 | transition metal ion binding |
| 12. | WP_172417888 | GO:0046914 | 0.97262 | transition metal ion binding |
| 13. | WP_172417888 | GO:0008270 | 0.96917 | zinc ion binding |
| 14. | WP_172418100 | GO:0046914 | 0.96635 | transition metal ion binding |
| 15. | WP_229844002 | GO:0046873 | 0.89965 | metal ion transmembrane transporter activity |
| 16. | WP_088249605 | GO:0008270 | 0.8993 | zinc ion binding |
| 17. | WP_143342227 | GO:0022857 | 0.87958 | transmembrane transporter activity |
| 18. | WP_143342227 | GO:0005215 | 0.87628 | Transporter activity |
| 19. | WP_143342227 | GO:0015075 | 0.83174 | ion transmembrane transporter activity |
| 17. | WP_088250461 | GO:0020037 | 0.83002 | heme binding |
| 18. | WP_088250474 | GO:0046914 | 0.82648 | transition metal ion binding |
| 19. | WP_088247726 | GO:0008270 | 0.82147 | zinc ion binding |
| 20. | WP_088247726 | GO:0046914 | 0.78883 | transition metal ion binding |
| 21. | WP_172418076 | GO:0046914 | 0.71328 | transition metal ion binding |
| 22. | WP_143342227 | GO:0015318 | 0.70012 | inorganic molecular entity<br>transmembrane transporter<br>activity |
| 23. | WP_172418079 | GO:0008270 | 0.69894 | zinc ion binding |
| 24. | WP_143341863 | GO:0046914 | 0.68499 | transition metal ion binding |
| 25. | WP_088250474 | GO:0005506 | 0.6793 | iron ion binding |
| 26. | WP_143342227 | GO:0042626 | 0.64938 | ATPase-coupled<br>transmembrane transporter<br>activity |
| 27. | WP_172418076 | GO:0030145 | 0.63205 | manganese ion binding |
| 28. | WP_172418079 | GO:0050897 | 0.63059 | cobalt ion binding |
| 29. | WP_088246697 | GO:0046914 | 0.62478 | transition metal ion binding |
| 30. | WP_143342227 | GO:0008324 | 0.6191 | cation transmembrane<br>transporter activity |
| 31. | WP_143342227 | GO:0022890 | 0.611 | inorganic cation transmembrane<br>transporter activity |
| 32. | WP_143342227 | GO:0015399 | 0.60611 | primary active transmembrane<br>transporter activity |
| 33. | WP_143342227 | GO:0022804 | 0.58785 | active transmembrane<br>transporter activity |
| 34. | WP_172418079 | GO:0046914 | 0.56284 | transition metal ion binding |
| 35. | WP_172418079 | GO:0008237 | 0.56269 | metallopeptidase activity |
| 36. | WP_088248339 | GO:0046914 | 0.55885 | transition metal ion binding |
| 37. | WP_143342227 | GO:0097367 | 0.54295 | carbohydrate derivative binding |
| 38. | WP_088248899 | GO:0051537 | 0.5484 | 2 iron, 2 sulfur cluster binding |
| 39. | WP_088250155 | GO:0008270 | 0.53643 | zinc ion binding |
| 40. | WP_143342227 | GO:0042625 | 0.53347 | ATPase-coupled ion<br>transmembrane transporter<br>activity |
| 41. | WP_143342227 | GO:0022853 | 0.5137 | active ion transmembrane transporter activity |
| 42. | WP_143342015 | GO:0051536 | 0.51247 | iron-sulfur cluster binding |
| 43. | WP_143342015 | GO:0051540 | 0.50994 | metal cluster binding |

**Figure 6:**
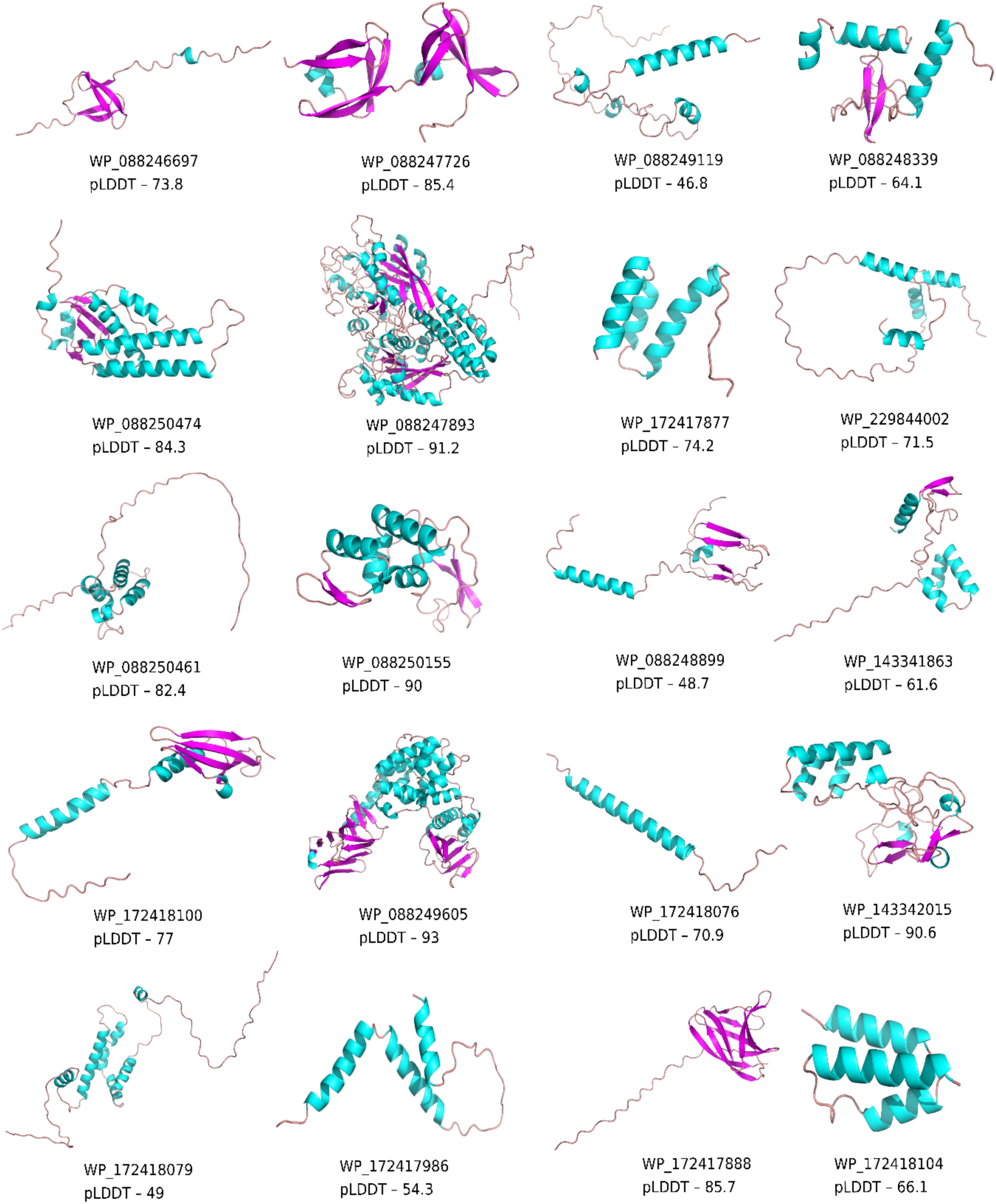
Images of all the 20 protein structures that have a metal ion binding property of 0.5 and above. The 20 protein structures that have been predicted to get bind with metals in *D. indicus* DR1, predicted by DeepFRI.

DeepFRI also predicts multifunctionality. Among the 1017 hypothetical proteins, 78 score ≥ 0.50 for at least one metal-binding term while simultaneously exhibiting other features such as ATPase activity, transporter function, transmembrane localization, or efflux-pump signatures. The counts rise to 92 and 115 at the 0.40 and 0.30 cutoffs, respectively (Table 4). GO term definitions were cross-checked against the Gene Ontology resource (Gene Ontology Consortium, et al., 2023)

**Table 4:**
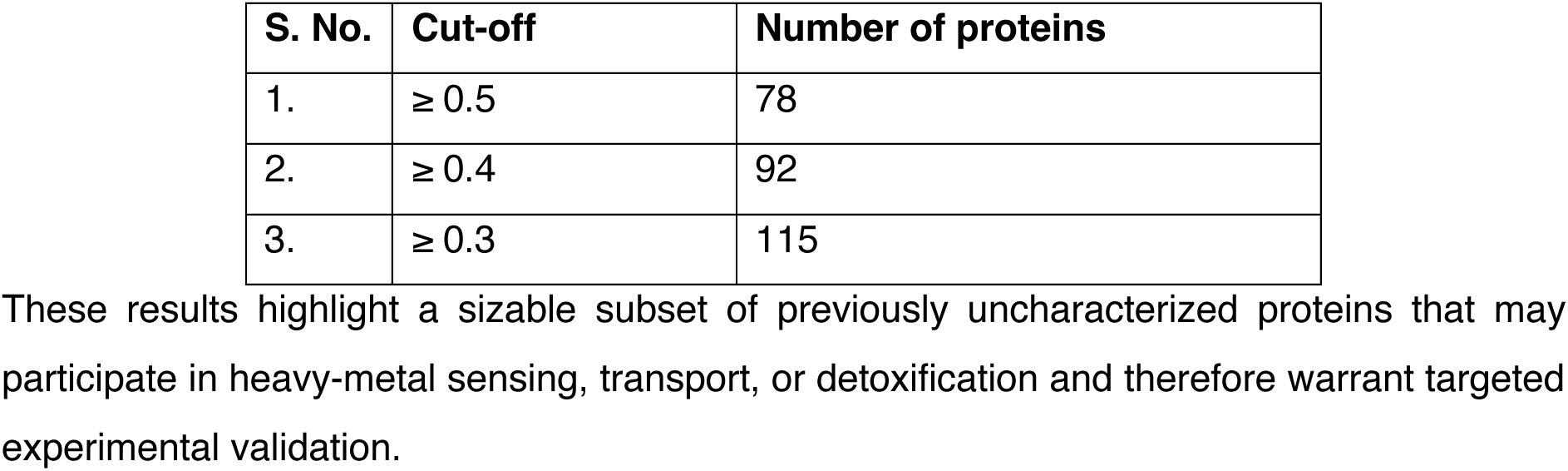
Number of proteins that are predicted by DeepFRI to possess metal ion binding property and other associated properties. It represents the number of proteins identified by DeepFRI as likely to possess metal ion binding properties and other associated properties.

## 4. Discussion

This study provides the first proteome-wide structural and functional annotation of *Deinococcus indicus* DR1. We modeled all 4128 proteins with AlphaFold2 and assigned CATH domains to 2735 of them; 1017 remained “hypothetical” at the outset. We then combined three orthogonal pipelines—DeepFRI, MorphologFinder (Foldseek + EggNOG-Mapper), and GenBank annotations—to explore each protein’s potential role in metal tolerance. The comparison reveals both convergence and notable discrepancies that highlight gaps in current databases.

### Chromium

Chromium resistance offers a representative case. Four proteins— WP_088249364, WP_088248628, WP_088247682, and WP_229844026—receive chromium-related annotations from MorphologFinder and GenBank (Supplementary Table 11), yet DeepFRI predicts none. All four are transporter-like, but whether they export chromium or merely share motifs with other metal efflux pumps is unclear. Because transition-metal transporters are often promiscuous, experimental assays will be required to confirm substrate range.

### Cobalt

A similar picture emerges for cobalt. GenBank and MorphologFinder converge on a single cobalamin-biosynthetic enzyme, WP_244898023 (Supplementary Table 12), whereas DeepFRI assigns cobalt-binding activity to two additional proteins, WP_172418079 and WP_172417973 (Supplementary Table 13), that lack EggNOG hits, underscoring how structure-based learning can uncover candidates missed by homology search.

### Copper

Copper exemplifies the challenge: GenBank annotates six copper-binding proteins-WP_088248749, WP_088250286, WP_088250320, WP_229844292, WP_229844418 and WP_244898031 (Table 5). DeepFRI flags eight copper-binding proteins-WP_088250185, WP_088249177, WP_088249243, WP_143342227, WP_143342125, WP_088247977, WP_143342094 and WP_216360397 (Table 6) but, in the absence of EggNOG support, these calls rest solely on the graph network. Foldseek offers alternative functions, leaving their actual role open. Resolving these discrepancies will require metal-binding or transport assays.

**Table 5:**
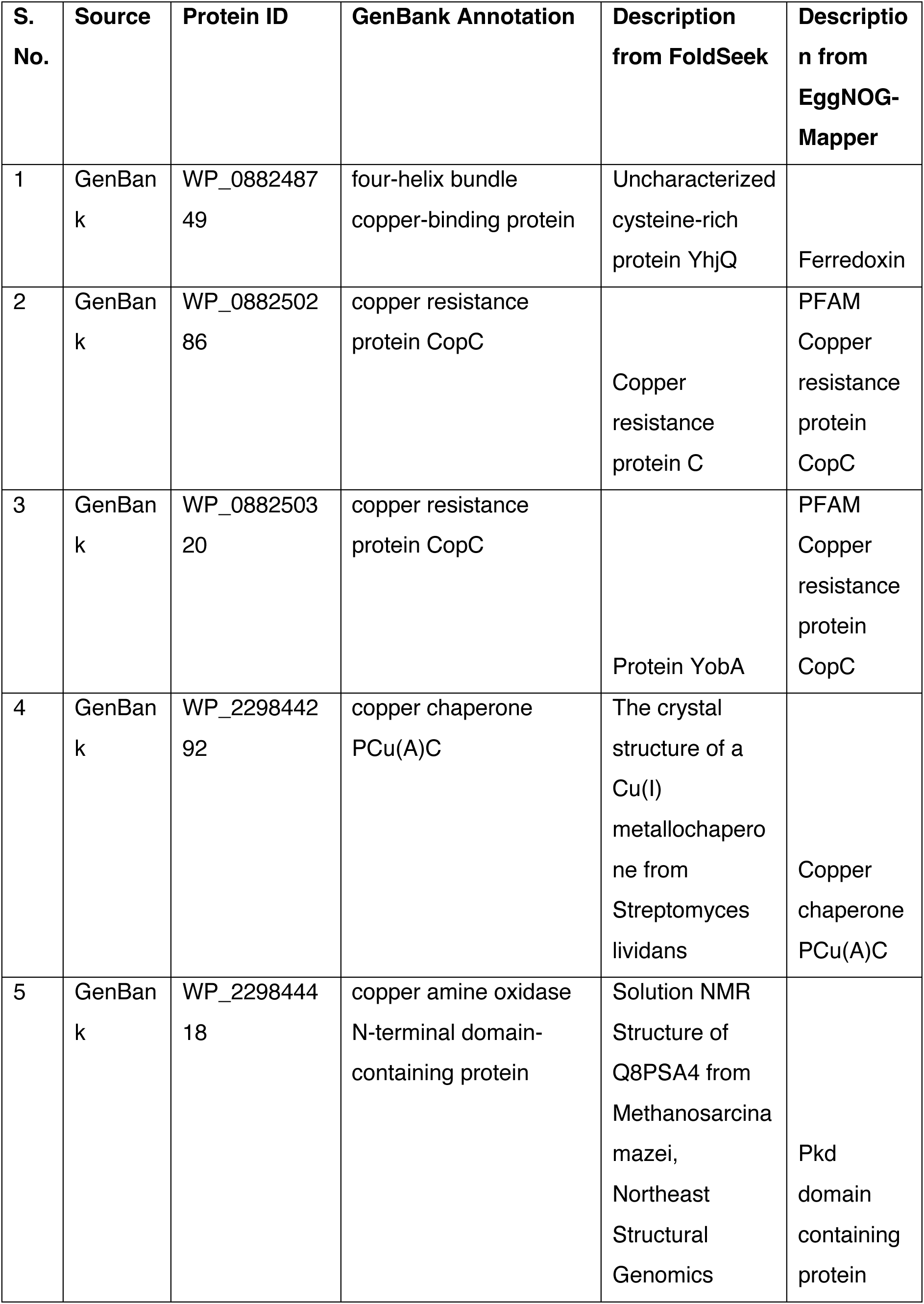
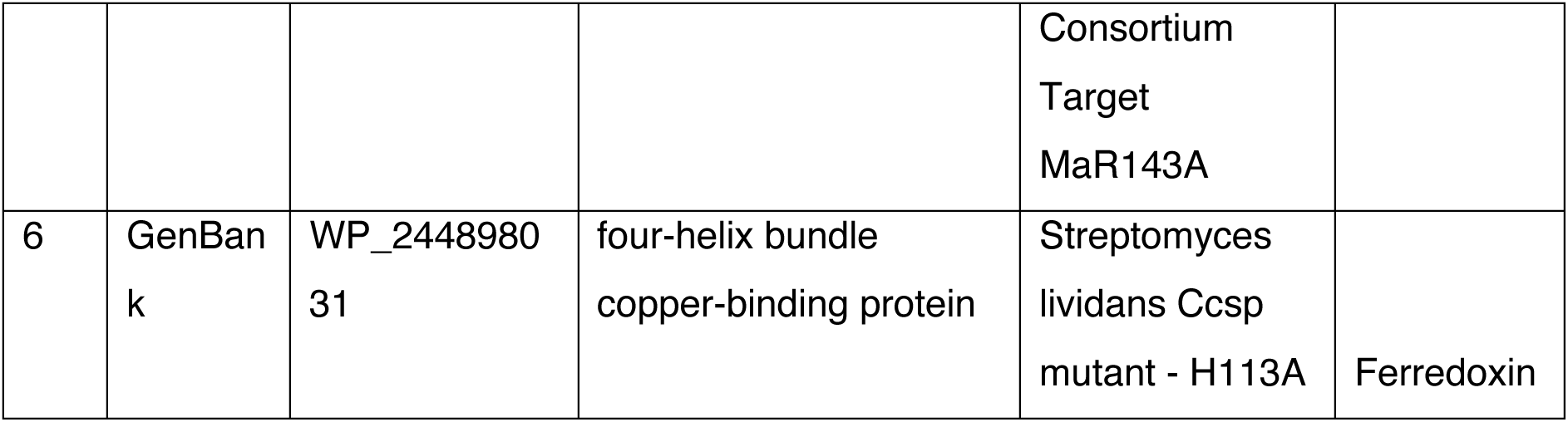
List of proteins that bind to copper as per GenBank annotation. It represents the list of proteins annotated in GenBank as having the ability to bind to copper and descriptive annotations obtained from FoldSeek and EggNOG-Mapper.

| S. No. | Source | Protein ID | GenBank Annotation | Description from FoldSeek | Description from EggNOG-Mapper |
| --- | --- | --- | --- | --- | --- |
| 1 | GenBank | WP_088248749 | four-helix bundle copper-binding protein | Uncharacterized cysteine-rich protein YhjQ | Ferredoxin |
| 2 | GenBank | WP_088250286 | copper resistance protein CopC | Copper resistance protein C | PFAM Copper resistance protein CopC |
| 3 | GenBank | WP_088250320 | copper resistance protein CopC | Protein YobA | PFAM Copper resistance protein CopC |
| 4 | GenBank | WP_229844292 | copper chaperone PCu(A)C | The crystal structure of a Cu(I) metallochaperone from *Streptomyces lividans* | Copper chaperone PCu(A)C |
| 5 | GenBank | WP_229844418 | copper amine oxidase N-terminal domain-containing protein | Solution NMR Structure of Q8PSA4 from *Methanosarcina mazei*, Northeast Structural Genomics | Pkd domain containing protein |
|  |  |  |  | Consortium<br>Target<br>MaR143A |  |
| 6 | GenBank | WP_244898031 | four-helix bundle<br>copper-binding protein | Streptomyces<br>lividans Ccsp<br>mutant - H113A | Ferredoxin |

**Table 6:**
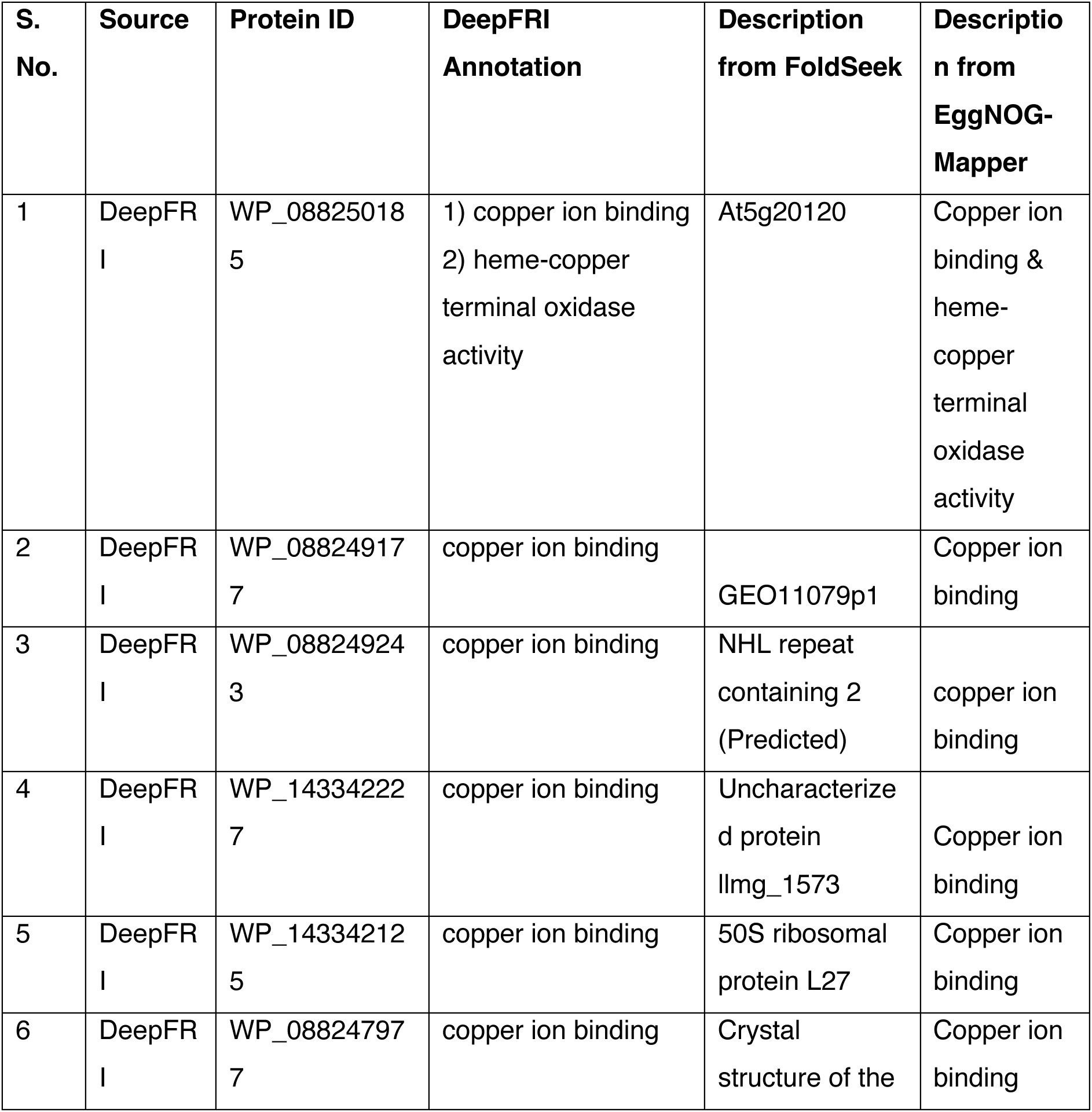
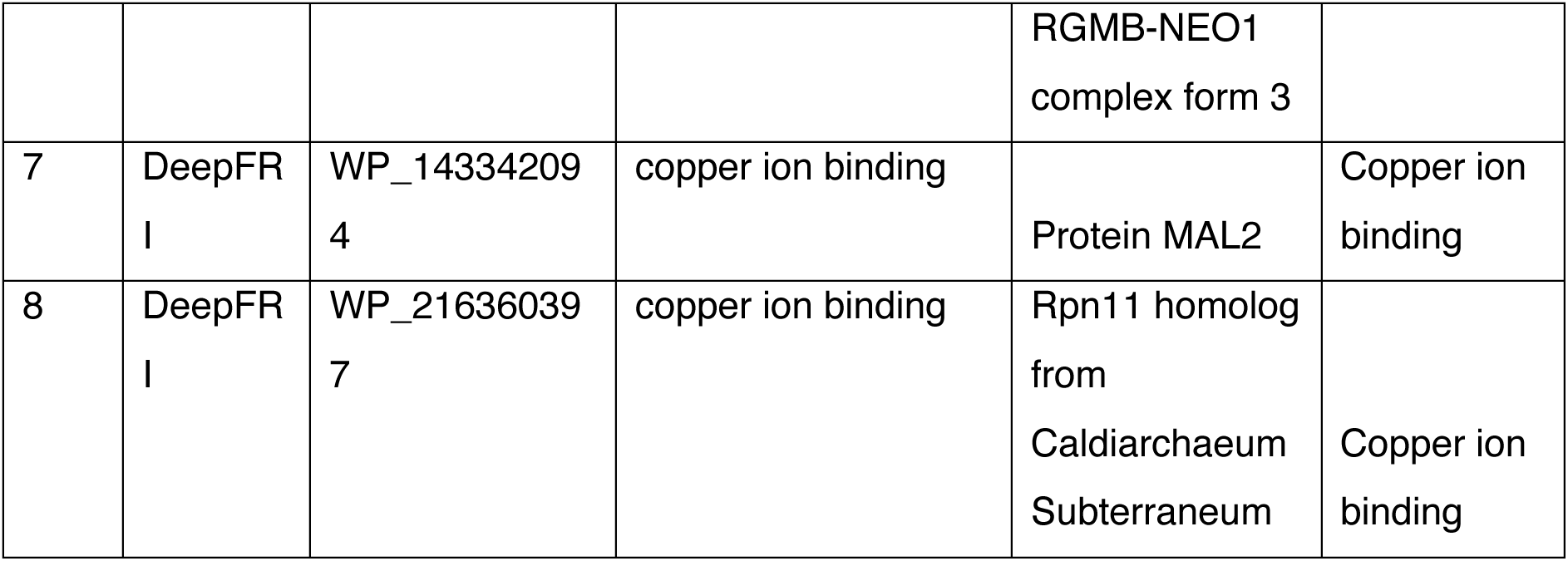
List of proteins that bind to copper as per DeepFRI annotation. It represents the list of proteins annotated in DeepFRI as having the ability to bind to copper and descriptive annotations obtained from FoldSeek and EggNOG-Mapper.

### Iron

For iron, GenBank lists forty-one proteins, twenty-seven of which are also detected by DeepFRI; only a subset agrees at the level of specific transporter or enzyme class, although WP_088246955 is consistently annotated as an iron ABC-transporter permease (Supplementary Tables 14 and 15).

### Manganese and Molybdenum

GenBank annotates two manganese-binding proteins, WP_088249680 and WP_088249842 (Supplementary Table 16). The former shows mismatched functions between GenBank and MorphologFinder, whereas the latter matches across both sources. DeepFRI adds four manganese candidates (WP_172418076, WP_088248334, WP_088246619, and WP_143341924; Supplementary Table 17), none of which receive EggNOG hits. For molybdenum, GenBank provides four candidates, with partial agreement from MorphologFinder (Supplementary Table 18).

### Nickel and Zinc

GenBank flags five nickel-related proteins, including WP_088248123 and WP_088248124, which align with MorphologFinder annotations, and WP_229844207 and WP_229844208, which lack EggNOG hits (Supplementary Table 19). DeepFRI predicts eleven nickel binders (Supplementary Table 20), but none match MorphologFinder, reflecting substantial functional ambiguity. Zinc shows a similar pattern: GenBank lists 28 proteins (Supplementary Table 21) and DeepFRI adds 26 more (Supplementary Table 22), yet overlap is limited and many entries lack EggNOG corroboration.

Taken together, these comparisons repeat the theme: overlapping but non-identical protein sets, with many instances where one pipeline produces no hit and another assigns a plausible but unverified function (Supplementary Tables 16–22).

Several patterns nevertheless support the idea that *D. indicus* DR1 deploys a broad metal-homeostasis toolkit. First, high-confidence DeepFRI predictions cluster around ABC-transporter permeases, P-type ATPases, and metal-binding oxidoreductases—machineries known to mediate detoxification in other bacteria. Second, CATH domain analysis shows a pronounced enrichment of nucleotide-binding folds, redox domains, and sensor modules (CHASE, PAS, GAF) that often regulate metal response. Third, many multidomain proteins combine these motifs—for example, the CHASE–PAS–GAF sensor WP_088247513 fused to kinase or phosphatase domains—implying integrated signaling circuits that coordinate arsenic reduction with oxidative-stress pathways.

Discrepancies among annotation pipelines likely stem from methodological biases. DeepFRI draws on structural graphs and thus excels when AlphaFold2 produces confident models but can misclassify highly flexible or novel folds. MorphologFinder, anchored in Foldseek structural alignment and EggNOG orthology, is conservative and may overlook distant analogs. GenBank entries inherit historical annotations that sometimes lack experimental support. Reconciling these views demands targeted experiments: metal-binding assays, growth tests under metal stress, and, where feasible, gene-knockout studies.

## 5. Conclusion

This structural-genomic survey of *Deinococcus indicus* DR1 shows how AI modeling can rapidly illuminate the biology of a non-model extremophile. AlphaFold2 produced “very-high”-confidence models (pLDDT ≥ 90) for 2,145 of the strain’s 4,128 proteins, turning more than half of the proteome from structure-dark to atomically mapped. CATH/InterPro analysis revealed ten highly recurrent domain types—dominated by nucleotide-binding, DNA-interaction, redox, and signaling folds—and highlighted a 12-domain RNA-polymerase β subunit plus six seven-domain enzymes that likely coordinate multi-step reactions.

Functionally, the combined DeepFRI and MorphologFinder pipelines assign plausible metal-binding or metal-transport roles to over 100 previously hypothetical proteins, spanning arsenic, chromium, cobalt, copper, iron, manganese, molybdenum, nickel, and zinc. This breadth suggests that *D. indicus* DR1 possesses a far more versatile metallome-management network.

The resulting atlas converts raw genome data into an actionable experimental roadmap. Twenty top-scoring candidate metal-binding proteins now form a priority list for expression, mutagenesis, and knockout studies aimed at confirming their roles in sequestration or efflux. Recurrent P-loop NTPase, Rossmann-fold, and GNAT motifs point to catalytic scaffolds that could be engineered into synthetic-biology circuits for bioremediation or bio-mining. All models, domain calls, and an interactive browser (https://deinococcus.in) are publicly available, offering ready-made hypotheses to the community. More broadly, the work illustrates how AI-driven structural genomics can accelerate functional discovery in extremophiles and lays a foundation for developing robust microbial platforms to mitigate heavy-metal pollution.

## Acknowledgements

The authors acknowledge SASTRA Deemed to be University for infrastructural support. The authors also thank Mr. Shashank Ravichandran for the website related help and support. RP acknowledges Shiv Nadar Institution of Eminence for infrastructural support. DS was supported by doctoral fellowship from Shiv Nadar Institution of Eminence.

## Data Availability

The modeled structures and all the metal functional data are available at https://deinococcus.in.

## CReDIT Statement

SDR curated the data, developed the website and wrote the manuscript. GV, SS, DS, and MT curated, processed, and compiled the data, and also edited the manuscript. RP and RMY conceptualized the idea, executed and approved the final version of the manuscript.

## References

Anantharaman, V., Aravind, L., 2001. The CHASE domain: a predicted ligand-binding module in plant cytokinin receptors and other eukaryotic and bacterial receptors. Trends Biochem Sci 26, 579–582. 10.1016/S0968-0004(01)01968-5

Aziz, M.F., Caetano-Anollés, G., 2021. Evolution of networks of protein domain organization. Sci Rep 11, 12075. 10.1038/s41598-021-90498-8

Barrera, A., Alastruey-Izquierdo, A., Martín, M.J., Cuesta, I., Vizcaíno, J.A., 2014. Analysis of the Protein Domain and Domain Architecture Content in Fungi and Its Application in the Search of New Antifungal Targets. PLoS Comput Biol 10, e1003733. 10.1371/journal.pcbi.1003733

Basu MK, Poliakov E, Rogozin IB. Domain mobility in proteins: functional and evolutionary implications. Brief Bioinform. 2009 May;10(3):205–16. doi: 10.1093/bib/bbn057

Bharat Siva Varma, P., Adimulam, Y.B., Kodukula, S., 2015. In silico functional annotation of a hypothetical protein from *Staphylococcus aureus*. J Infect Public Health 8, 526–532. 10.1016/j.jiph.2015.03.007

Bryant P. Deep learning for protein complex structure prediction. Curr Opin Struct Biol. 2023 Apr;79:102529. doi: 10.1016/j.sbi.2023.102529

Cavaco, L.M., Hasman, H., Stegger, M., Andersen, P.S., Skov, R., Fluit, A.C., Ito, T., Aarestrup, F.M., 2010. Cloning and Occurrence of czrC, a Gene Conferring Cadmium and Zinc Resistance in Methicillin-Resistant *Staphylococcus aureus* CC398 Isolates. Antimicrob Agents Chemother 54, 3605–3608. 10.1128/AAC.00058-10

Chauhan, D., Srivastava, P.A., Agnihotri, V., Yennamalli, R.M., Priyadarshini, R., 2019. Structure and function prediction of arsenate reductase from *Deinococcus indicus* DR1. J Mol Model 25, 15. 10.1007/s00894-018-3885-3

Chauhan, D., Srivastava, P.A., Yennamalli, R.M., Priyadarshini, R., 2017. Draft Genome Sequence of *Deinococcus indicus* DR1, a Novel Strain Isolated from a Freshwater Wetland. Genome Announc 5. 10.1128/genomeA.00754-17

Dhanapal, A.R., Venkidasamy, B., Solai Ramatchandirane, P., 2021. Molecular characterization of stress tolerance genes associated with *D. indicus* strain under extreme environment conditions. Environ Geochem Health 43, 4905–4917. 10.1007/s10653-020-00788-9

Eggers, S., Safdar, N., Malecki, K.M., 2018. Heavy metal exposure and nasal *Staphylococcus aureus* colonization: analysis of the National Health and Nutrition Examination Survey (NHANES). Environmental Health 17, 2. 10.1186/s12940-017-0349-7

Gene Ontology Consortium; Aleksander SA, Balhoff J, Carbon S, Cherry JM, Drabkin HJ, Ebert D, Feuermann M, Gaudet P, Harris NL, Hill DP, Lee R, Mi H, Moxon S, Mungall CJ, Muruganugan A, Mushayahama T, Sternberg PW, Thomas PD, Van Auken K, Ramsey J, Siegele DA, Chisholm RL, Fey P, Aspromonte MC, Nugnes MV, Quaglia F, Tosatto S, Giglio M, Nadendla S, Antonazzo G, Attrill H, Dos Santos G, Marygold S, Strelets V, Tabone CJ, Thurmond J, Zhou P, Ahmed SH, Asanitthong P, Luna Buitrago D, Erdol MN, Gage MC, Ali Kadhum M, Li KYC, Long M, Michalak A, Pesala A, Pritazahra A, Saverimuttu SCC, Su R, Thurlow KE, Lovering RC, Logie C, Oliferenko S, Blake J, Christie K, Corbani L, Dolan ME, Drabkin HJ, Hill DP, Ni L, Sitnikov D, Smith C, Cuzick A, Seager J, Cooper L, Elser J, Jaiswal P, Gupta P, Jaiswal P, Naithani S, Lera-Ramirez M, Rutherford K, Wood V, De Pons JL, Dwinell MR, Hayman GT, Kaldunski ML, Kwitek AE, Laulederkind SJF, Tutaj MA, Vedi M, Wang SJ, D’Eustachio P, Aimo L, Axelsen K, Bridge A, Hyka-Nouspikel N, Morgat A, Aleksander SA, Cherry JM, Engel SR, Karra K, Miyasato SR, Nash RS, Skrzypek MS, Weng S, Wong ED, Bakker E, Berardini TZ, Reiser L, Auchincloss A, Axelsen K, Argoud-Puy G, Blatter MC, Boutet E, Breuza L, Bridge A, Casals-Casas C, Coudert E, Estreicher A, Livia Famiglietti M, Feuermann M, Gos A, Gruaz-Gumowski N, Hulo C, Hyka-Nouspikel N, Jungo F, Le Mercier P, Lieberherr D, Masson P, Morgat A, Pedruzzi I, Pourcel L, Poux S, Rivoire C, Sundaram S, Bateman A, Bowler-Barnett E, Bye-A-Jee H, Denny P, Ignatchenko A, Ishtiaq R, Lock A, Lussi Y, Magrane M, Martin MJ, Orchard S, Raposo P, Speretta E, Tyagi N, Warner K, Zaru R, Diehl AD, Lee R, Chan J, Diamantakis S, Raciti D, Zarowiecki M, Fisher M, James-Zorn C, Ponferrada V, Zorn A, Ramachandran S, Ruzicka L, Westerfield M. The Gene Ontology knowledgebase in 2023. Genetics. 2023 May 4;224(1):iyad031. doi: 10.1093/genetics/iyad031

Gerber, E., Bernard, R., Castang, S., Chabot, N., Coze, F., Dreux-Zigha, A., Hauser, E., Hivin, P., Joseph, P., Lazarelli, C., Letellier, G., Olive, J., Leonetti, J. -P., 2015. Deinococcus as new chassis for industrial biotechnology: biology, physiology and tools. J Appl Microbiol 119, 1–10. 10.1111/jam.12808

Ghosh, S., Osman, S., Vaishampayan, P., Venkateswaran, K., 2010. Recurrent Isolation of Extremotolerant Bacteria from the Clean Room Where Phoenix Spacecraft Components Were Assembled. Astrobiology 10, 325–335. 10.1089/ast.2009.0396

Gligorijević, V., Renfrew, P.D., Kosciolek, T., Leman, J.K., Berenberg, D., Vatanen, T., Chandler, C., Taylor, B.C., Fisk, I.M., Vlamakis, H., Xavier, R.J., Knight, R., Cho, K., Bonneau, R., 2021. Structure-based protein function prediction using graph convolutional networks. Nat Commun 12, 3168. 10.1038/s41467-021-23303-9

Igiri BE, Okoduwa SIR, Idoko GO, Akabuogu EP, Adeyi AO, Ejiogu IK. Toxicity and Bioremediation of Heavy Metals Contaminated Ecosystem from Tannery Wastewater: A Review. J Toxicol. 2018 Sep 27;2018:2568038. doi: 10.1155/2018/2568038

Ijaq, J., Malik, G., Kumar, A., Das, P.S., Meena, N., Bethi, N., Sundararajan, V.S., Suravajhala, P., 2019. A model to predict the function of hypothetical proteins through a nine-point classification scoring schema. BMC Bioinformatics 20, 14. 10.1186/s12859-018-2554-y

Jones, P., Binns, D., Chang, H.-Y., Fraser, M., Li, W., McAnulla, C., McWilliam, H., Maslen, J., Mitchell, A., Nuka, G., Pesseat, S., Quinn, A.F., Sangrador-Vegas, A., Scheremetjew, M., Yong, S.-Y., Lopez, R., Hunter, S., 2014. InterProScan 5: genome-scale protein function classification. Bioinformatics 30, 1236–1240. 10.1093/bioinformatics/btu031

Jumper, J., Evans, R., Pritzel, A., Green, T., Figurnov, M., Ronneberger, O., Tunyasuvunakool, K., Bates, R., Žídek, A., Potapenko, A., Bridgland, A., Meyer, C., Kohl, S.A.A., Ballard, A.J., Cowie, A., Romera-Paredes, B., Nikolov, S., Jain, R., Adler, J., Back, T., Petersen, S., Reiman, D., Clancy, E., Zielinski, M., Steinegger, M., Pacholska, M., Berghammer, T., Bodenstein, S., Silver, D., Vinyals, O., Senior, A.W., Kavukcuoglu, K., Kohli, P., Hassabis, D., 2021. Highly accurate protein structure prediction with AlphaFold. Nature 596, 583–589. 10.1038/s41586-021-03819-2

Möglich, A., Ayers, R.A., Moffat, K., 2009. Structure and Signaling Mechanism of Per-ARNT-Sim Domains. Structure 17, 1282–1294. 10.1016/j.str.2009.08.011

Paysan-Lafosse, T., Blum, M., Chuguransky, S., Grego, T., Pinto, B.L., Salazar, G.A., Bileschi, M.L., Bork, P., Bridge, A., Colwell, L., Gough, J., Haft, D.H., Letunić, I., Marchler-Bauer, A., Mi, H., Natale, D.A., Orengo, C.A., Pandurangan, A.P., Rivoire, C., Sigrist, C.J.A., Sillitoe, I., Thanki, N., Thomas, P.D., Tosatto, S.C.E., Wu, C.H., Bateman, A., 2023. InterPro in 2022. Nucleic Acids Res 51, D418–D427. 10.1093/nar/gkac993

Jaafar, R., Al-Sulami, A., Al-Taee, A., Aldoghachi, F., Suhaimi, N., and Mohammed, S. 2016. Biosorption of some Heavy Metals by *Deinococcus radiodurans* Isolated from Soil in Basra Governorate-Iraq. J Bioremediat Biodegrad 07. 10.4172/2155-6199.1000332

Ranganathan, S., Sethi, D., Kasivisweswaran, S., Ramya, L., Priyadarshini, R., Yennamalli, R.M., 2023. Structural and functional mapping of ars gene cluster in *Deinococcus indicus* DR1. Comput Struct Biotechnol J 21, 519–534. 10.1016/j.csbj.2022.12.015

Sharma, R., Rensing, C., Rosen, B.P., Mitra, B., 2000. The ATP Hydrolytic Activity of Purified ZntA, a Pb(II)/Cd(II)/Zn(II)-translocating ATPase from *Escherichia coli*. Journal of Biological Chemistry 275, 3873–3878. 10.1074/jbc.275.6.3873

Sillitoe, I., Bordin, N., Dawson, N., Waman, V.P., Ashford, P., Scholes, H.M., Pang, C.S.M., Woodridge, L., Rauer, C., Sen, N., Abbasian, M., Le Cornu, S., Lam, S.D., Berka, K., Varekova, I.H., Svobodova, R., Lees, J., Orengo, C.A., 2021. CATH: increased structural coverage of functional space. Nucleic Acids Res 49, D266–D273. 10.1093/nar/gkaa1079

Suresh, K., Reddy, G.S.N., Sengupta, S., Shivaji, S., 2004. *Deinococcus indicus* sp. nov., an arsenic-resistant bacterium from an aquifer in West Bengal, India. Int J Syst Evol Microbiol 54, 457–461. 10.1099/ijs.0.02758-0

Tsai, K.J., Yoon, K.P., Lynn, A.R., 1992. ATP-dependent cadmium transport by the cadA cadmium resistance determinant in everted membrane vesicles of *Bacillus subtilis*. J Bacteriol 174, 116–121. 10.1128/jb.174.1.116-121.1992

Varadi, M., Anyango, S., Deshpande, M., Nair, S., Natassia, C., Yordanova, G., Yuan, D., Stroe, O., Wood, G., Laydon, A., Žídek, A., Green, T., Tunyasuvunakool, K., Petersen, S., Jumper, J., Clancy, E., Green, R., Vora, A., Lutfi, M., Figurnov, M., Cowie, A., Hobbs, N., Kohli, P., Kleywegt, G., Birney, E., Hassabis, D., Velankar, S., 2022. AlphaFold Protein Structure Database: massively expanding the structural coverage of protein-sequence space with high-accuracy models. Nucleic Acids Res 50, D439–D444. 10.1093/nar/gkab1061

Wang Y, Zhang H, Zhong H, Xue Z. Protein domain identification methods and online resources. Comput Struct Biotechnol J. 2021 Feb 2;19:1145–1153. doi: 10.1016/j.csbj.2021.01.041

Weon, H.-Y., Kim, B.-Y., Schumann, P., Son, J.-A., Jang, J., Go, S.-J., Kwon, S.-W., 2007. *Deinococcus cellulosilyticus* sp. nov., isolated from air. Int J Syst Evol Microbiol 57, 1685– 1688. 10.1099/ijs.0.64951-0

