## Supplementary Tables for "Mapping the Metalloproteome of *Deinococcus indicus* DR1 through Integrative Structure and Function Annotation"

**Supplementary Table 1: pLDDT score distribution throughout the proteome of *D. indicus* DR1.** It represents the number of proteins that fall under every pLDDT score category.

| **pLDDT score range** | **No. of proteins** |
| --- | --- |
| 90 and above | 2175 |
| 70 – 89 | 1688 |
| 50 – 69 | 229 |
| 49 and lower | 66 |

**Supplementary Table 2: Number of proteins and the domain count.** It represents the number of proteins having each particular domain count.

| **S. No.** | **Domain Count** | **No. of Proteins** |
| --- | --- | --- |
| 1. | 1 | 1553 |
| 2. | 2 | 775 |
| 3. | 3 | 258 |
| 4. | 4 | 81 |
| 5. | 5 | 44 |
| 6. | 6 | 17 |
| 7. | 7 | 6 |
| 8. | 12 | 1 |

**Supplementary Table 3: List of CATH domains, its region and its function of the twelve-domain protein WP_088248407.** It represents the comprehensive list detailing the CATH domains identified in the 12-domain protein WP_088248407, including the specific regions each domain occupies within the protein sequence, along with a description of function associated with each domain.

| **S. no.** | **Region** | | **CATH ID** | **Domain name** |
| --- | --- | --- | --- | --- |
|  | **From** | **To** |  |  |
| 1. | 1 | 124 | 4.10.860.120 | RNA polymerase II, clamp domain |
| 2. | 147 | 207 | 2.40.50.100 | RNA polymerase II |
| 3. | 247 | 320 | 2.40.50.100 | RNA polymerase II |
| 4. | 411 | 468 | 2.40.50.100 | RNA polymerase II |
| 5. | 624 | 785 | 2.40.40.20 | DNA-directed RNA polymerase subunit |
| 6. | 661 | 705 | 1.10.40.90 | DNA-directed RNA polymerase subunit beta |
| 7. | 786 | 964 | 1.10.274.100 | RNA polymerase Rpb1, domain 3 |
| 8. | 965 | 1102 | 1.10.132.30 | RNA polymerase Rpb1, funnel domain |
| 9. | 1129 | 1201 | 3.90.105.10 | Molybdopretin biosynthesis moea protein, domain 2 |
| 10 | 1283 | 1348 | 2.40.50.100 | RNA polymerase II |
| 11. | 1267 | 1446 | 1.10.1790.20 | DNA-directed RNA polymerase subunit beta |
| 12. | 1488 | 1511 | 1.10.150.390 | DNA-directed RNA polymerase subunit |

**Supplementary Table 4: List of CATH domains, its region and its function of the seven-domain protein WP_088247437.** It represents the comprehensive list detailing the CATH domains identified in the 7-domain protein WP_088247437, including the specific regions each domain occupies within the protein sequence, along with a description of function associated with each domain.

| **S.no.** | **Region** | | **CATH ID** | **Domain name** |
| --- | --- | --- | --- | --- |
|  | **From** | **To** |  |  |
| 1. | 16 | 122 | 2.60.40.1110 | Not yet named |
| 2. | 152 | 257 | 2.60.40.1110 | Not yet named |
| 3. | 259 | 364 | 2.60.40.1110 | Not yet named |
| 4. | 375 | 490 | 2.60.40.1130 | Rab geranylgeranyltransferase alpha-subunit, insert domain |
| 5. | 491 | 600 | 2.60.40.10 | Immunoglobulins |
| 6. | 601 | 1164 | 3.20.20.80 | Glycosidases |
| 7. | 1165 | 1233 | 2.60.40.1180 | Golgi alpha-mannosidase II |

**Supplementary Table 5: List of CATH domains, its region and its function of the seven-domain protein WP_088247513.** It represents the comprehensive list detailing the CATH domains identified in the seven-domain protein WP_088247513, including the specific regions each domain occupies within the protein sequence, along with a description of function associated with each domain.

| **S no.** | **Region** | | **CATH ID** | **Domain name** |
| --- | --- | --- | --- | --- |
|  | **From** | **To** |  |  |
| 1. | 78 | 256 | 3.30.450.350 | CHASE domain |
| 2. | 329 | 452 | 3.30.450.20 | PAS domain |
| 3. | 453 | 574 | 3.30.450.20 | PAS domain |
| 4. | 575 | 684 | 3.30.450.20 | PAS domain |
| 5. | 719 | 850 | 3.30.450.40 | GAF domian |
| 6. | 860 | 946 | 1.10.287.130 | RNA-binding domian |
| 7. | 949 | 1100 | 3.30.565.10 | Histidine kinase-like ATPase |

**Supplementary Table 6: List of CATH domains, its region and its function of the seven-domain protein WP_088247559**. It represents the comprehensive list detailing the CATH domains identified in the seven-domain protein WP_088247559, including the specific regions each domain occupies within the protein sequence, along with a description of function associated with each domain.

| **S no.** | **Region** | | **CATH ID** | **Domain name** |
| --- | --- | --- | --- | --- |
|  | **From** | **To** |  |  |
| 1. | 1 | 73 | 1.10.287.610 | Helix hairpin bin |
| 2. | 74 | 324 | 3.30.470.30 | DNA ligase/mRNA capping enzyme |
| 3. | 326 | 399 | 2.40.50.140 | Nucleic acid binding protein |
| 4. | 401 | 437 | 6.20.10.30 | Not yet named |
| 5. | 438 | 512 | 1.10.150.20 | 5' to 3' exonuclease, C-terminal subdomain |
| 6. | 513 | 592 | 1.10.150.20 | 5' to 3' exonuclease, C-terminal subdomain |
| 7. | 595 | 678 | 3.40.50.10190 | BRCT domain |

**Supplementary Table 7: List of CATH domains, its region and its function of the seven-domain protein WP_088248081.** It represents the comprehensive list detailing the CATH domains identified in the seven-domain protein WP_088248081, including the specific regions each domain occupies within the protein sequence, along with a description of function associated with each domain.

| **S. no.** | **Region** | | **CATH ID** | **Domain name** |
| --- | --- | --- | --- | --- |
|  | **From** | **To** |  |  |
| 1. | 12 | 114 | 3.30.450.20 | PAS domain |
| 2. | 115 | 276 | 3.30.450.40 | GAF domain |
| 3. | 288 | 444 | 3.30.450.40 | GAF domain |
| 4. | 457 | 631 | 3.30.450.40 | GAF domain |
| 5. | 632 | 803 | 3.30.450.40 | GAF domain |
| 6. | 804 | 884 | 1.10.287.130 | Signal transduction histidine kinase, dimerisation/phosphotransfer (DHp) domain. |
| 7. | 885 | 1034 | 3.30.565.10 | Histidine kinase-like ATPase |

**Supplementary Table 8: List of CATH domains, its region and its function of the seven-domain protein WP_088248246.** It represents the comprehensive list detailing the CATH domains identified in the 7-domain protein WP_088248246, including the specific regions each domain occupies within the protein sequence, along with a description of function associated with each domain.

| **S. no.** | **Region** | | **CATH ID** | **Domain name** |
| --- | --- | --- | --- | --- |
|  | **From** | **To** |  |  |
| 1. | 1 | 147 | 3.10.20.740 | Not yet named |
| 2. | 156 | 228 | 3.30.70.20 | Not yet named |
| 3. | 241 | 294 | 2.20.25.90 | ADC-like domains |
| 4. | 306 | 367 | 3.40.50.740 | Not yet named |
| 5. | 368 | 408 | 3.40.228.10 | Dimethylsulfoxide Reductase |
| 6. | 469 | 531 | 3.40.50.1220 | TPP-binding domain |
| 7. | 552 | 676 | 3.40.50.740 | Not yet named |

**Supplementary Table 9: List of CATH domains, its region and its function of the seven-domain protein WP_088248452.** It represents the comprehensive list detailing the CATH domains identified in the 7-domain protein WP_088248452, including the specific regions each domain occupies within the protein sequence, along with a description of function associated with each domain.

| **S. no.** | **Region** | | **CATH ID** | **Domain name** |
| --- | --- | --- | --- | --- |
|  | **From** | **To** |  |  |
| 1. | 13 | 1036 | 2.40.270.10 | DNA-directed RNA polymerase, subunit 2, domain 6 |
| 2. | 16 | 616 | 3.90.1100.10 | Not yet named |
| 3. | 154 | 347 | 3.90.1100.10 | Not yet named |
| 4. | 493 | 559 | 2.30.150.10 | DNA-directed RNA polymerase, beta subunit, external 1 domain |
| 5. | 617 | 700 | 2.40.50.100 | RNA polymerase II/Efflux pump adaptor protein, barrel-sandwich hybrid domain |
| 6. | 736 | 863 | 2.40.50.150 | RNA polymerase II, Rpb2 subunit, wall domain |
| 7. | 1046 | 1095 | 3.90.1800.10 | RNA polymerase alpha subunit dimerisation domain |

**Supplementary Table 10: Number of proteins that are predicted by DeepFRI to possess metal ion binding property.** It represents the number of proteins identified by DeepFRI as likely to possess metal ion binding properties.

| **S. No.** | **Cut-off** | **Number of proteins** |
| --- | --- | --- |
| 1. | ≥ 0.5 | 20 |
| 2. | ≥ 0.4 | 27 |
| 3. | ≥ 0.3 | 35 |

**Supplementary Table 11: List of proteins that bind to Chromium as per GenBank annotation.** It represents the list of proteins annotated in GenBank as having the ability to bind to chromium and descriptive annotations obtained from FoldSeek and EggNOG-Mapper.

| **S.No.** | **Source** | **Protein ID** | **GenBank Annotation** | **Description from Foldseek** | **Description from**  **eggNOG-Mapper** |
| --- | --- | --- | --- | --- | --- |
| 1 | GenBank | WP_088249364 | chromate transporter | Probable transporter | Chromate transporter |
| 2 | GenBank | WP_088248628 | chromate transporter | Probable transporter | Chromate transporter |
| 3 | GenBank | WP_088247682 | chromate transporter | Uncharacterized transporter YwrA | Chromate transporter |
| 4 | GenBank | WP_229844026 | chromate transporter | Uncharacterized transporter YwrB | Chromate transporter |

**Supplementary Table 12: List of proteins that bind to Cobalt as per GenBank annotation.** It represents the list of proteins annotated in GenBank as having the ability to bind to cobalt and descriptive annotations obtained from FoldSeek and EggNOG-Mapper.

| **S. No.** | **Source** | **Protein ID** | **GenBank Annotation** | **Description from Foldseek** | **Description from**  **eggNOG-Mapper** |
| --- | --- | --- | --- | --- | --- |
| 1 | GenBank | WP_244898023 | cobaltochelatase subunit CobN | Cobaltochelatase, CobN subunit | TIGRFAM cobaltochelatase, CobN subunit |

**Supplementary Table 13: List of proteins that bind to Cobalt as per DeepFRI annotation.** It represents the list of proteins annotated in DeepFRI as having the ability to bind to cobalt and descriptive annotations obtained from FoldSeek and EggNOG-Mapper.

| **S. No.** | **Source** | **Protein ID** | **DeepFRI Annotation** | **Description from Foldseek** | **Description from**  **eggNOG-Mapper** |
| --- | --- | --- | --- | --- | --- |
| 1 | DeepFri | WP_172418079 | cobalt ion binding | Cobalt ion binding | Cobalt ion binding |
| 2 | DeepFri | WP_172417973 | cobalt ion binding | Putative ABC-type multidrug transport system, ATPase component | Cobalt ion binding |

**Supplementary Table 14: List of proteins that bind to Iron as per GenBank annotation.** It represents the list of proteins annotated in GenBank as having the ability to bind to iron and descriptive annotations obtained from FoldSeek and EggNOG-Mapper.

| **S. No.** | **Source** | **Protein ID** | **GenBank Annotation** | **Description from FoldSeek** | **Description from**  **EggNOG-Mapper** |
| --- | --- | --- | --- | --- | --- |
| 1 | GenBank | WP_055363595 | Fe-S cluster assembly ATPase SufC | ABC transporter domain-containing protein | COG0396 ABC-type transport system involved in Fe-S cluster assembly ATPase component |
| 2 | GenBank | WP_088246955 | iron ABC transporter permease | Iron (III)-transport system permease HitB | ABC-type Fe3 transport system permease component |
| 3 | GenBank | WP_088247038 | SUF system NifU family Fe-S cluster assembly protein | Crystal structure of Iron-sulfur cluster biosynthesis protein IscU (TTHA1736) from thermus thermophilus HB8 | SUF system FeS assembly protein |
| 4 | GenBank | WP_088247105 | iron ABC transporter permease | ABC transporter permease | PFAM Binding-protein-dependent transport system inner membrane component |
| 5 | GenBank | WP_088247310 | (2Fe-2S)-binding protein | Nicotinate dehydrogenase subunit A | COG2080 Aerobic-type carbon monoxide dehydrogenase, small subunit CoxS CutS homologs |
| 6 | GenBank | WP_088247317 | (2Fe-2S)-binding protein | Carbon monoxide dehydrogenase small chain | COG2080 Aerobic-type carbon monoxide dehydrogenase, small subunit CoxS CutS homologs |
| 7 | GenBank | WP_088247333 | iron-siderophore ABC transporter substrate-binding protein | Putative ABC transporter substrate-binding lipoprotein YhfQ | abc-type fe3 -hydroxamate transport system, periplasmic component |
| 8 | GenBank | WP_088247401 | L-serine ammonia-lyase, iron-sulfur-dependent subunit beta | L-serine deaminase | TIGRFAM L-serine dehydratase, iron-sulfur-dependent, beta subunit |
| 9 | GenBank | WP_088247456 | heterodisulfide reductase-related iron-sulfur binding cluster | Crystal structure of a putative Asp/Glu Racemase from Yersinia pestis | Fe-S oxidoreductase |
| 10 | GenBank | WP_088247735 | iron-sulfur cluster binding | Uncharacterized protein MJ0966 | PFAM Radical SAM |
| 11 | GenBank | WP_088247751 | quinolinate synthase NadA | Quinolinate synthase A | Catalyzes the condensation of iminoaspartate with dihydroxyacetone phosphate to form quinolinate |
| 12 | GenBank | WP_088247907 | L-serine ammonia-lyase, iron-sulfur-dependent, subunit alpha | Probable L-serine dehydratase, alpha chain | TIGRFAM L-serine dehydratase, iron-sulfur-dependent, alpha subunit |
| 13 | GenBank | WP_088247988 | iron-siderophore ABC transporter substrate-binding protein | Fe(3+)-citrate-binding protein YfmC | abc-type fe3 -hydroxamate transport system, periplasmic component |
| 14 | GenBank | WP_088247991 | iron ABC transporter permease | Iron-regulated surface determinant protein F | Belongs to the binding-protein-dependent transport system permease family. FecCD subfamily |
| 15 | GenBank | WP_088248030 | 2Fe-2S iron-sulfur cluster-binding protein | Aldehyde oxidoreductase iron-sulfur-binding subunit PaoA | COG2080 Aerobic-type carbon monoxide dehydrogenase, small subunit CoxS CutS homologs |
| 16 | GenBank | WP_088248138 | iron-siderophore ABC transporter substrate-binding protein | Petrobactin-binding protein FpuA | COG0614 ABC-type Fe3 -hydroxamate transport system, periplasmic component |
| 17 | GenBank | WP_088248143 | (2Fe-2S)-binding protein | Crystal structure of an aldehyde oxidase from Methylobacillus sp. KY4400 | COG2080 Aerobic-type carbon monoxide dehydrogenase, small subunit CoxS CutS homologs |
| 18 | GenBank | WP_088248163 | GTP 3',8-cyclase MoaA | GTP 3',8-cyclase | Catalyzes the cyclization of GTP to (8S)-3',8-cyclo-7,8- dihydroguanosine 5'-triphosphate |
| 19 | GenBank | WP_088248179 | Fe-S cluster assembly protein SufD | UPF0051 protein SSP1857 | TIGRFAM FeS assembly protein SufD |
| 20 | GenBank | WP_088248181 | Fe-S cluster assembly protein SufB | UPF0051 protein SE_0610 | ABC-type transport system involved in Fe-S cluster assembly, permease component |
| 21 | GenBank | WP_088248610 | Dps family protein | DNA protection during starvation protein 1 | Belongs to the Dps family |
| 22 | GenBank | WP_088249062 | nitrite/sulfite reductase | Sulfite reductase [ferredoxin] | Nitrite and sulphite reductase 4Fe-4S |
| 23 | GenBank | WP_088249078 | glycolate oxidase subunit GlcF | Probable glycolate oxidase iron-sulfur subunit | 4Fe-4S dicluster domain |
| 24 | GenBank | WP_088249142 | tRNA (N6-isopentenyl adenosine(37)-C2)-methylthiotransferase MiaB | tRNA-2-methylthio-N(6)-dimethylallyladenosine synthase | Catalyzes the methylthiolation of N6- (dimethylallyl)adenosine (i(6)A), leading to the formation of 2- methylthio-N6-(dimethylallyl)adenosine (ms(2)i(6)A) at position 37 in tRNAs that read codons beginning with uridine |
| 25 | GenBank | WP_088249411 | iron ABC transporter permease | Inward-facing conformation of ABC heme importer BhuUV from Burkholderia cenocepacia | Belongs to the binding-protein-dependent transport system permease family. FecCD subfamily |
| 26 | GenBank | WP_088249702 | (2Fe-2S)-binding protein | Crystal structure of glyceraldehyde oxidoreductase | 2 iron, 2 sulfur cluster binding |
| 27 | GenBank | WP_088249716 | 4 iron, 4 sulfur cluster binding | L-lysine 2,3-aminomutase | TIGRFAM KamA family protein |
| 28 | GenBank | WP_088249880 | succinate dehydrogenase iron-sulfur subunit | Succinate dehydrogenase iron-sulfur subunit | TIGRFAM succinate dehydrogenase and fumarate reductase iron-sulfur protein |
| 29 | GenBank | WP_088249957 | 4 iron, 4 sulfur cluster binding | Epoxyqueuosine reductase | TIGRFAM iron-sulfur cluster binding protein |
| 30 | GenBank | WP_088250006 | 4 iron, 4 sulfur cluster binding | Lipoyl synthase | Catalyzes the radical-mediated insertion of two sulfur atoms into the C-6 and C-8 positions of the octanoyl moiety bound to the lipoyl domains of lipoate-dependent enzymes, thereby converting the octanoylated domains into lipoylated derivatives |
| 31 | GenBank | WP_088250411 | Rieske 2Fe-2S domain-containing protein | Crystal Structure of the Rieske protein from Thermus thermophilus | Catalyzes the radical-mediated insertion of two sulfur atoms into the C-6 and C-8 positions of the octanoyl moiety bound to the lipoyl domains of lipoate-dependent enzymes, thereby converting the octanoylated domains into lipoylated derivatives |
| 32 | GenBank | WP_088250414 | iron-sulfur cluster assembly accessory protein | iron-sulfur cluster binding | Catalyzes the synthesis of the hydroxymethylpyrimidine phosphate (HMP-P) moiety of thiamine from aminoimidazole ribotide (AIR) in a radical S-adenosyl-L-methionine (SAM)-dependent reaction |
| 33 | GenBank | WP_088250468 | iron-sulfur cluster binding | Phosphomethylpyrimidine synthase | Catalyzes the synthesis of the hydroxymethylpyrimidine phosphate (HMP-P) moiety of thiamine from aminoimidazole ribotide (AIR) in a radical S-adenosyl-L-methionine (SAM)-dependent reaction |
| 34 | GenBank | WP_172417882 | 2Fe-2S iron-sulfur cluster-binding protein | Na(+)-translocating NADH-quinone reductase subunit F | PFAM 2Fe-2S iron-sulfur cluster binding domain |
| 35 | GenBank | WP_172418032 | 4 iron, 4 sulfur cluster binding | Ribosomal protein S12 methylthiotransferase RimO | Catalyzes the methylthiolation of an aspartic acid residue of ribosomal protein S12 |
| 36 | GenBank | WP_172418047 | iron ABC transporter permease | Transporter | Belongs to the binding-protein-dependent transport system permease family. FecCD subfamily |
| 37 | GenBank | WP_229844051 | 4 iron, 4 sulfur cluster | Type-4 uracil-DNA glycosylase | TIGRFAM Phage SPO1 DNA polymerase-related protein |
| 38 | GenBank | WP_229844259 | ubiquinol-cytochrome c reductase iron-sulfur subunit | Crystal structure of arsenite oxidase from Alcaligenes faecalis (Af Aio) bound to antimony oxyanion | PFAM Rieske 2Fe-2S domain |
| 39 | GenBank | WP_229844332 | non-heme iron oxygenase ferredoxin subunit | 3-phenylpropionate/cinnamic acid dioxygenase ferredoxin subunit | COGs COG2146 Ferredoxin subunits of nitrite reductase and ring-hydroxylating dioxygenase |
| 40 | GenBank | WP_229844372 | 4 iron, 4 sulfur cluster binding | Crystal structure of Deinococcus radiodurans Endonuclease III-1 R61Q variant | DNA repair enzyme that has both DNA N-glycosylase activity and AP-lyase activity. The DNA N-glycosylase activity releases various damaged pyrimidines from DNA by cleaving the N- glycosidic bond, leaving an AP (apurinic apyrimidinic) site. The AP-lyase activity cleaves the phosphodiester bond 3' to the AP site by a beta-elimination, leaving a 3'-terminal unsaturated sugar and a product with a terminal 5'-phosphate |
| 41 | GenBank | WP_244898005 | iron ABC transporter permease | Fe(3+)-citrate import system permease protein YfmE | Belongs to the binding-protein-dependent transport system permease family. FecCD subfamily |

**Supplementary Table 15: List of proteins that bind to Iron as per DeepFRI annotation.** It represents the list of proteins annotated in DeepFRI as having the ability to bind to iron and descriptive annotations obtained from FoldSeek and EggNOG-Mapper.

| **S. No.** | **Source** | **Protein ID** | **DeepFRI Annotation** | **Description from FoldSeek** | **Description from**  **EggNOG-Mapper** |
| --- | --- | --- | --- | --- | --- |
| 1 | DeepFRI | WP_172417986 | 1. iron-sulfur cluster binding 2. 4 iron, 4 sulfur cluster binding | Uncharacterized protein | iron-sulfur cluster binding & 4 iron, 4 sulfur cluster binding |
| 2 | DeepFRI | WP_172417877 | 1) iron-sulfur cluster binding 2) 2 iron, 2 sulfur cluster binding 3) iron ion binding | Methionine--tRNA ligase, cytoplasmic | iron-sulfur cluster binding & 2 iron, 2 sulfur cluster binding |
| 3 | DeepFRI | WP_088250474 | iron ion binding | Methionine--tRNA ligase, cytoplasmic | iron ion binding |
| 4 | DeepFRI | WP_088248899 | 1) 2 iron, 2 sulfur cluster binding 2) iron-sulfur cluster binding | Si:dkey-111e8.4 | 1) 2 iron, 2 sulfur cluster binding 2) iron-sulfur cluster binding |
| 5 | DeepFRI | WP_143342015 | 1) iron-sulfur cluster binding 2) 2 iron, 2 sulfur cluster binding 3) 3 iron, 4 sulfur cluster binding 4) iron ion binding | 2.25 Angstrom Resolution Crystal Structure of 6-phospho-alpha-glucosidase from Klebsiella pneumoniae in Complex with NAD. | 1) iron-sulfur cluster binding 2) 2 iron, 2 sulfur cluster binding 3) 3 iron, 4 sulfur cluster binding 4) Iron ion binding |
| 6 | DeepFRI | WP_143342125 | 1) iron-sulfur cluster binding 2) 2 iron, 2 sulfur cluster binding 3) 4 iron, 4 sulfur cluster binding | 50S ribosomal protein L27 | 1) iron-sulfur cluster binding 2) 2 iron, 2 sulfur cluster binding 3) 4 iron, 4 sulfur cluster binding |
| 7 | DeepFRI | WP_172418104 | iron ion binding | IL7R in complex with an antagonist | iron ion binding |
| 8 | DeepFRI | WP_088246697 | iron-sulfur cluster binding | ArgX from Sulfolobus tokodaii complexed with LysW/Glu/ADP/Mg/Zn/Sulfate | iron-sulfur cluster binding |
| 9 | DeepFRI | WP_143341863 | iron-sulfur cluster binding | Uncharacterized protein | iron-sulfur cluster binding |
| 10 | DeepFRI | WP_088248607 | iron-sulfur cluster binding | Uncharacterized protein | iron-sulfur cluster binding |
| 11 | DeepFRI | WP_088250408 | 1) iron-sulfur cluster binding 2) 2 iron, 2 sulfur cluster binding | Crystal structure of Greglin in complex with subtilisin | 1) ron-sulfur cluster binding 2) 2 iron, 2 sulfur cluster binding |
| 12 | DeepFRI | WP_088247001 | 1) 4 iron, 4 sulfur cluster binding 2) iron ion binding 3) iron-sulphur binding | Crystal structure of the P95883_SULSO protein from Sulfolobus solfataricus. NESG target SsR10. | 1) 4 iron, 4 sulfur cluster binding 2) iron ion binding 3) iron-sulphur binding |
| 13 | DeepFRI | WP_088248044 | iron ion binding | Nitroreductase domain-containing protein | iron ion binding |
| 14 | DeepFRI | WP_088248991 | 1) 2 iron, 2 sulfur cluster binding 2) iron-sulfur cluster binding | Crystal structure of ProN-Tk-SP from Thermococcus kodakaraensis | 1) 2 iron, 2 sulfur cluster binding 2) iron-sulfur cluster binding |
| 15 | DeepFRI | WP_088246912 | iron ion binding | B-block binding subunit of TFIIIC | iron ion binding |
| 16 | DeepFRI | WP_088248226 | iron ion binding | Uncharacterized protein | Protein conserved in bacteria |
| 17 | DeepFRI | WP_088247460 | 1) iron-sulfur cluster binding 2) iron ion binding | Uncharacterized protein | 1) iron-sulfur cluster binding 2) iron ion binding |
| 18 | DeepFRI | WP_088246616 | 1) iron-sulfur cluster binding 2) 2 iron, 2 sulfur cluster binding | The crystal structure of lobe domain of E. coli RNA polymerase complexed with the C-terminal domain of UvrD | 1) Iron-sulphur cluster binding 2) 2 iron, 2 sulfur cluster binding |
| 19 | DeepFRI | WP_088247436 | 1) iron-sulfur cluster binding 2) iron ion binding | Synaptic vesicle glycoprotein 2A | iron-sulfur cluster binding & iron ion binding |
| 20 | DeepFRI | WP_216360397 | 1) iron-sulfur cluster binding 2) iron ion binding | Rpn11 homolog from Caldiarchaeum Subterraneum | 1) iron-sulfur cluster binding   2) iron ion binding |
| 21 | DeepFRI | WP_143342094 | iron-sulfur cluster binding | Protein MAL2 | iron-sulfur cluster binding |
| 22 | DeepFRI | WP_088248631 | iron-sulfur cluster binding | Cytochrome b mRNA maturase bI3 | iron-sulfur cluster binding |
| 23 | DeepFRI | WP_088247460 | 1) 4 iron, 4 sulfur cluster binding 2) 2 iron, 2 sulfur cluster binding | Uncharacterized protein | 1) 4 iron, 4 sulfur cluster binding   2 iron, 2 sulfur cluster binding |
| 24 | DeepFRI | WP_088246741 | iron-sulfur cluster binding | HTH-type transcriptional regulator BhcR | iron-sulfur cluster binding |
| 25 | DeepFRI | WP_088247977 | 2 iron, 2 sulfur cluster binding | Crystal structure of the RGMB-NEO1 complex form 3 | 2 iron, 2 sulfur cluster binding |
| 26 | DeepFRI | WP_088248339 | iron-sulfur cluster binding | Probable chromosome-partitioning protein ParB | iron-sulfur cluster binding |
| 27 | DeepFRI | WP_088250147 | 2 iron, 2 sulfur cluster binding | Uncharacterized protein | 2 iron, 2 sulfur cluster binding |

**Supplementary Table 16: List of proteins that bind to Manganese as per GenBank annotation.** It represents the list of proteins annotated in GenBank as having the ability to bind to manganese and descriptive annotations obtained from FoldSeek and EggNOG-Mapper.

| **S. No.** | **Source** | **Protein ID** | **GenBank Annotation** | **Description from FoldSeek** | **Description from**  **EggNOG-Mapper** |
| --- | --- | --- | --- | --- | --- |
| 1 | GenBank | WP_088249680 | manganese ion binding | The structure of a CoA pyrophosphatase from D. Radiodurans | PFAM NUDIX hydrolase |
| 2 | GenBank | WP_088249842 | manganese-dependent inorganic pyrophosphatase | Probable manganese-dependent inorganic pyrophosphatase | inorganic diphosphatase activity |

**Supplementary Table 17: List of proteins that bind to Manganese as per DeepFRI annotation.** It represents the list of proteins annotated in DeepFRI as having the ability to bind to manganese and descriptive annotations obtained from FoldSeek and EggNOG-Mapper.

| **S. No.** | **Source** | **Protein ID** | **DeepFRI Annotation** | **Description from FoldSeek** | **Description from**  **EggNOG-Mapper** |
| --- | --- | --- | --- | --- | --- |
| 1 | DeepFRI | WP_172418076 | manganese ion binding | ZP domain-containing protein | manganese ion binding |
| 2 | DeepFRI | WP_088248334 | manganese ion binding | Mediator of RNA polymerase II transcription subunit 11 | manganese ion binding |
| 3 | DeepFRI | WP_088246619 | manganese ion binding | Structure of mouse CXorf40A, Selenomethionine derivative | manganese ion binding |
| 4 | DeepFRI | WP_143341924 | manganese ion binding | Uncharacterized protein | manganese ion binding |

**Supplementary Table 18: List of proteins that bind to Molybdenum as per GenBank annotation.** It represents the list of proteins annotated in GenBank as having the ability to bind to molybdenum and descriptive annotations obtained from FoldSeek and EggNOG-Mapper.

| **S. No.** | **Source** | **Protein ID** | **GenBank Annotation** | **Description from FoldSeek** | **Description from**  **EggNOG-Mapper** |
| --- | --- | --- | --- | --- | --- |
| 1 | GenBank | WP_088247211 | molybdopterin molybdotransferase MoeA | Molybdopterin molybdenumtransferase | MoeA N-terminal region (domain I and II) |
| 2 | GenBank | WP_088247448 | molybdenum cofactor biosynthesis protein B | Molybdenum cofactor biosynthesis protein B | Molybdenum cofactor biosynthesis protein B |
| 3 | GenBank | WP_229844140 | molybdenum cofactor guanylyltransferase | Probable molybdenum cofactor guanylyltransferase | Transfers a GMP moiety from GTP to Mo-molybdopterin (Mo- MPT) cofactor (Moco or molybdenum cofactor) to form Mo- molybdopterin guanine dinucleotide (Mo-MGD) cofactor |
| 4 | GenBank | WP_229844368 | molybdenum ion binding | MOSC domain-containing protein | PFAM MOSC domain |

**Supplementary Table 19: List of proteins that bind to Nickel as per GenBank annotation.** It represents the list of proteins annotated in GenBank as having the ability to bind to nickel and descriptive annotations obtained from FoldSeek and EggNOG-Mapper.

| **S. No.** | **Source** | **Protein ID** | **GenBank Annotation** | **Description from FoldSeek** | **Description from**  **EggNOG-Mapper** |
| --- | --- | --- | --- | --- | --- |
| 1 | GenBank | WP_088248121 | nickel cation binding | Urease subunit alpha | Belongs to the metallo-dependent hydrolases superfamily. Urease alpha subunit family |
| 2 | GenBank | WP_088248123 | hydrogenase maturation nickel HypA metallochaperone | Hydrogenase maturation factor HypA | Hydrogenase/urease nickel incorporation, metallochaperone, hypA |
| 3 | GenBank | WP_088248124 | hydrogenase nickel incorporation protein HypB | Hydrogenase maturation factor HypB | CobW/HypB/UreG, nucleotide-binding domain |
| 4 | GenBank | WP_229844207 | nickel cation binding | Urease accessory protein UreD | nickel cation binding |
| 5 | GenBank | WP_229844208 | nickel cation binding | Urease accessory protein UreF | nickel cation binding |

**Supplementary Table 20: List of proteins that bind to Nickel as per DeepFRI annotation** It represents the list of proteins annotated in DeepFRI as having the ability to bind to nickel and descriptive annotations obtained from FoldSeek and EggNOG-Mapper.

| **S. No.** | **Source** | **Protein ID** | **DeepFRI Annotation** | **Description from FoldSeek** | **Description from**  **EggNOG-Mapper** |
| --- | --- | --- | --- | --- | --- |
| 1 | DeepFRI | WP_088248231 | nickel cation binding | Thioredoxin | cell redox homeostasis |
| 2 | DeepFRI | WP_244898015 | nickel cation binding | Molybdopterin synthase sulfur carrier subunit | nickel cation binding |
| 3 | DeepFRI | WP_088247726 | nickel cation binding | Alpha-aminoadipate carrier protein LysW | Transposase, IS605 OrfB family |
| 4 | DeepFRI | WP_088246875 | nickel cation binding | Crystal structure of predicted HD superfamily hydrolase (104161995) from uncultured Thermotogales bacterium at 1.45 A resolution | mRNA catabolic process |
| 5 | DeepFRI | WP_088248228 | nickel cation binding | Crystal structure of domains AC3-AC5 of yeast acetyl-CoA carboxylase | nickel cation binding |
| 6 | DeepFRI | WP_088246741 | nickel cation binding | HTH-type transcriptional regulator BhcR | nickel cation binding |
| 7 | DeepFRI | WP_088249335 | nickel cation binding | Crystal structure of Ni-containing superoxide dismutase with Ni-ligation corresponding to the state after partial x-ray-induced reduction | nickel cation binding |
| 8 | DeepFRI | WP_229844093 | nickel cation binding | ATP-dependent zinc metalloprotease YME1L | nickel cation binding |
| 9 | DeepFRI | WP_088246585 | nickel cation binding | Crystal Structure of Drosophila melanogaster Pur-alpha | nickel cation binding |
| 10 | DeepFRI | WP_088248301 | nickel cation binding | Structure of the C-terminal non-repetitive domain of the spider dragline silk protein ADF-3 | nickel cation binding |
| 11 | DeepFRI | WP_088247666 | nickel cation binding | Uncharacterized protein | isomerase activity |

**Supplementary Table 21: List of proteins that bind to Zinc as per GenBank annotation.** It represents the list of proteins annotated in GenBank as having the ability to bind to zinc and descriptive annotations obtained from FoldSeek and EggNOG-Mapper.

| **S. No.** | **Source** | **Protein ID** | **GenBank Annotation** | **Description from FoldSeek** | **Description from**  **EggNOG-Mapper** |
| --- | --- | --- | --- | --- | --- |
| 1 | GenBank | WP_088246726 | zinc finger-like domain-containing protein | Crystal structure of Seabream Antiquitin and Elucidation of its substrate specificity | nickel cation binding |
| 2 | GenBank | WP_088246877 | zinc-dependent alcohol dehydrogenase family protein | Alcohol dehydrogenase GroES domain-containing protein | alcohol dehydrogenase |
| 3 | GenBank | WP_088246952 | SprT family zinc-dependent metalloprotease | Uncharacterized protein | Protein of unknown function DUF45 |
| 4 | GenBank | WP_088246953 | neutral zinc metallopeptidase | Uncharacterized protein YpfJ | Putative neutral zinc metallopeptidase |
| 5 | GenBank | WP_088247203 | zinc ABC transporter substrate-binding protein | ABC transporter substrate-binding protein | Belongs to the bacterial solute-binding protein 9 family |
| 6 | GenBank | WP_088247238 | zinc ion binding | CMP/dCMP-type deaminase domain-containing protein | MafB19-like deaminase |
| 7 | GenBank | WP_088247466 | zinc ion binding | Acrylyl-CoA reductase AcuI | COG0604 NADPH quinone reductase and related Zn-dependent |
| 8 | GenBank | WP_088247475 | zinc ion binding | Methionine synthase | Vitamin B12 dependent methionine synthase activation |
| 9 | GenBank | WP_088247776 | zinc metallopeptidase | Putative membrane protease YugP | PFAM peptidase, membrane zinc metallopeptidase |
| 10 | GenBank | WP_088248188 | zinc-dependent alcohol dehydrogenase | S-(hydroxymethyl)glutathione dehydrogenase | Alcohol dehydrogenase GroES-associated |
| 11 | GenBank | WP_088248659 | zinc ion binding | Crystal structure of M32 carboxypeptidase from Deinococcus radiodurans R1 | Broad specificity carboxypetidase that releases amino acids sequentially from the C-terminus, including neutral, aromatic, polar and basic residues |
| 12 | GenBank | WP_088248677 | zinc ion binding | PZ PEPTIDASE A with inhibitor 2 | metalloendopeptidase activity |
| 13 | GenBank | WP_088248709 | zinc ion binding | Serine hydroxymethyltransferase | Catalyzes the reversible interconversion of serine and glycine with tetrahydrofolate (THF) serving as the one-carbon carrier. This reaction serves as the major source of one-carbon groups required for the biosynthesis of purines, thymidylate, methionine, and other important biomolecules. Also exhibits THF- independent aldolase activity toward beta-hydroxyamino acids, producing glycine and aldehydes, via a retro-aldol mechanism |
| 14 | GenBank | WP_088248762 | zinc ion binding | Crystal structure of RNase J complexed with RNA | An RNase that has 5'-3' exonuclease and possibly endonuclease activity. Involved in maturation of rRNA and in some organisms also mRNA maturation and or decay |
| 15 | GenBank | WP_088248765 | zinc ion binding | ATP-dependent Clp protease ATP-binding subunit ClpX | ATP-dependent specificity component of the Clp protease. It directs the protease to specific substrates. Can perform chaperone functions in the absence of ClpP |
| 16 | GenBank | WP_088249459 | zinc ribbon domain-containing protein | IS200/IS605 family element transposase accessory protein TnpB |  |
| 17 | GenBank | WP_088249517 | zinc ribbon domain-containing protein | zf-RING_7 domain-containing protein | Zn-ribbon protein, possibly nucleic acid-binding |
| 18 | GenBank | WP_088249596 | SWIM zinc finger family protein | SWIM-type domain-containing protein | zinc ion binding |
| 19 | GenBank | WP_088249643 | M14 family zinc carboxypeptidase | Carboxypeptidase T | Zn_pept |
| 20 | GenBank | WP_088249762 | zinc ion binding | Probable bifunctional transcriptional activator/DNA repair enzyme AlkA | sequence-specific DNA binding |
| 21 | GenBank | WP_088249944 | zinc ion binding | M3 family oligoendopeptidase | Oligoendopeptidase, pepF M3 family |
| 22 | GenBank | WP_088250113 | ATP-dependent zinc metalloprotease FtsH | ATP-dependent zinc metalloprotease FtsH 2 | Acts as a processive, ATP-dependent zinc metallopeptidase for both cytoplasmic and membrane proteins. Plays a role in the quality control of integral membrane proteins |
| 23 | GenBank | WP_088250250 | ATP-dependent zinc metalloprotease FtsH | ATP-dependent zinc metalloprotease FtsH | Acts as a processive, ATP-dependent zinc metallopeptidase for both cytoplasmic and membrane proteins. Plays a role in the quality control of integral membrane proteins |
| 24 | GenBank | WP_088250491 | zinc-binding dehydrogenase | Zinc binding dehydrogenase-like protein | Zinc-binding alcohol dehydrogenase |
| 25 | GenBank | WP_088250512 | zinc ribbon domain-containing protein | Putative virulence protein | zinc ribbon domain-containing protein |
| 26 | GenBank | WP_172418180 | zinc ion binding | Cytosine deaminase | tRNA wobble adenosine to inosine editing |
| 27 | GenBank | WP_216360386 | zinc ion binding | Crystal structure of thermostable Carboxypeptidase (FisCP) from Fervidobacterium Islandicum AW-1 | Broad specificity carboxypetidase that releases amino acids sequentially from the C-terminus, including neutral, aromatic, polar and basic residues |
| 28 | GenBank | WP_244897980 | ATP-dependent zinc metalloprotease FtsH | ATP-dependent zinc metalloprotease FtsH | Acts as a processive, ATP-dependent zinc metallopeptidase for both cytoplasmic and membrane proteins. Plays a role in the quality control of integral membrane proteins |

**Supplementary Table 22: List of proteins that bind to Zinc as per DeepFRI annotation.** It represents the list of proteins annotated in DeepFRI as having the ability to bind to zinc and descriptive annotations obtained from FoldSeek and EggNOG-Mapper.

| **S. No.** | **Source** | **Protein ID** | **DeepFRI Annotation** | **Description from FoldSeek** | **Description from**  **EggNOG-Mapper** |
| --- | --- | --- | --- | --- | --- |
| 1 | DeepFRI | WP_172418100 | zinc ion binding | Ubiquitin-conjugating enzyme | zinc ion binding |
| 2 | DeepFRI | WP_172417888 | zinc ion binding | Putative periplasmic protein | zinc ion binding |
| 3 | DeepFRI | WP_088249605 | zinc ion binding | NMR structure of a designed cold unfolding four helix bundle | zinc ion binding |
| 4 | DeepFRI | WP_088247726 | zinc ion binding | Alpha-aminoadipate carrier protein LysW | zinc ion binding |
| 5 | DeepFRI | WP_172418079 | zinc ion binding | zinc ion binding | zinc ion binding |
| 6 | DeepFRI | WP_088250155 | zinc ion binding | Hydrogenase maturation factor HypA | zinc ion binding |
| 7 | DeepFRI | WP_088248339 | zinc ion binding | Probable chromosome-partitioning protein ParB | zinc ion binding |
| 8 | DeepFRI | WP_088246697 | zinc ion binding | ArgX from Sulfolobus tokodaii complexed with LysW/Glu/ADP/Mg/Zn/Sulfate | zinc ion binding |
| 9 | DeepFRI | WP_229844399 | zinc ion binding | DNA-directed RNA polymerase subunit | zinc ion binding |
| 10 | DeepFRI | WP_143341863 | zinc ion binding | Uncharacterized protein | zinc ion binding |
| 11 | DeepFRI | WP_143342058 | zinc ion binding | BTB domain-containing protein | zinc ion binding |
| 12 | DeepFRI | WP_088247190 | zinc ion binding | Os05g0156800 protein | zinc ion binding |
| 13 | DeepFRI | WP_216360387 | zinc ion binding | DNA topoisomerase 3 | zinc ion binding |
| 14 | DeepFRI | WP_172418004 | zinc ion binding | Crystal Structure of the Bacteriophage T4 recombination mediator protein UvsY, Lattice Type IV | zinc ion binding |
| 15 | DeepFRI | WP_088248607 | zinc ion binding | Os01g0615850 protein | zinc ion binding |
| 16 | DeepFRI | WP_088246704 | zinc ion binding | RhsD protein | zinc ion binding |
| 17 | DeepFRI | WP_088249408 | zinc ion binding | Crystal structure of a protein with unknown function from DUF155 family (YP_292156.1) from Prochlorococcus sp. NATL2A at 1.80 A resolution | zinc ion binding |
| 18 | DeepFRI | WP_088248337 | zinc ion binding | Translation machinery-associated protein 22 | zinc ion binding |
| 19 | DeepFRI | WP_088246585 | zinc ion binding | Crystal Structure of Drosophila melanogaster Pur-alpha | zinc ion binding |
| 20 | DeepFRI | WP_143342125 | zinc ion binding | 50S ribosomal protein L27 | zinc ion binding |
| 21 | DeepFRI | WP_143342015 | zinc ion binding | 2.25 Angstrom Resolution Crystal Structure of 6-phospho-alpha-glucosidase from Klebsiella pneumoniae in Complex with NAD | zinc ion binding |
| 22 | DeepFRI | WP_088247001 | zinc ion binding | Crystal structure of the P95883_SULSO protein from Sulfolobus solfataricus. NESG target SsR10. | zinc ion binding |
| 23 | DeepFRI | WP_143342202 | zinc ion binding | Bilin biosynthesis protein CpeZ | zinc ion binding |
| 24 | DeepFRI | WP_088247893 | zinc ion binding | Uncharacterized protein | zinc ion binding |
| 25 | DeepFRI | WP_088248044 | zinc ion binding | Nitroreductase domain-containing protein | zinc ion binding |
| 26 | DeepFRI | WP_088250063 | zinc ion binding | Oligogalacturonate lyase in complex with manganese | zinc ion binding |
